# Frequent, context-dependent effects of human genetic variation on Cas9 activity revealed by population-scale GUIDE-seq-2 and deep combinatorial CHANCE-seq profiling

**DOI:** 10.1101/2025.02.10.637517

**Authors:** Ashley R. Flory, Cicera R. Lazzarotto, Yichao Li, Jacqueline Chyr, Muyu Yang, Varun Katta, Elizabeth Urbina, GaHyun Lee, Rachael Wood, Azusa Matsubara, Claire Greene Whitfield, Sara R. Rashkin, Jian Ma, Yong Cheng, Shengdar Q. Tsai

## Abstract

Genome editing enzymes can introduce targeted changes to DNA in living cells^1–4^, transforming biological research and enabling the first approved gene editing therapy for sickle cell disease^5^. However, their activity can be altered by genetic variation at on- or off-target sites^6–8^, potentially impacting both their precision and therapeutic safety. Due to a lack of scalable methods to measure genome-wide editing activity in cells from large populations and diverse target libraries, the frequency and extent of these variant effects on editing remain unknown. Here, we present the first systematic, population-scale study of how genetic variation affects the cellular genome-wide activity of CRISPR-Cas9, enabled by a novel, sensitive, and unbiased cellular assay, GUIDE-seq-2, with improved scalability and accuracy compared to the original broadly adopted method^9^. Analyzing Cas9 genome-wide activity at 1,115 on- and off-target sites across six guide RNAs in cells from 95 individuals spanning four genetically diverse populations, we found that genetic variants frequently overlap off-target sites, with 14% significantly altering Cas9 editing activity. To understand the effect of mismatches in more diverse sequence contexts, we developed a novel method, combinatorial high-throughput analysis of nuclease cleavage effects (CHANCE-seq), the first massively parallel biochemical approach that can quantify Cas9 activity across millions of mismatched target sites. We leveraged this large-scale CHANCE-seq dataset to train a context-aware deep neural network model, CHANCE-net, to accurately predict and interpret the effects of single-nucleotide variants on off-targets with up to six mismatches. Our deep combinatorial profiling of Cas9 off-target activity revealed frequent, context-dependent synergistic effects of mismatches. Taken together, our findings illuminate an approach to accounting for genetic variation when designing genome-editing strategies for research and therapeutics.

---

Genome editing technology has potential to yield transformative therapies for a wide range of genetic diseases. The recent regulatory approval of exagamglogene autotemcel (exa-cel)—a Cas9 genome-edited autologous hematopoietic stem cell therapy that induces fetal hemoglobin to treat sickle cell disease—in the US and Europe^10,11^ marks a major milestone in this exciting field. Several other genome editing therapies are currently undergoing clinical trials^12^.

However, genome editors can also introduce unintended modifications at off-target sites^13^, such as large-scale chromosomal translocations or deletions associated with DSBs^9,14,15^. This unintended, off-target editing can potentially confound biological research and pose safety risks in therapeutic applications^16^. For example, Lorenzini et al. describe an off-target site in the intron of *USP9X* gene that resulted in disrupted expression of this gene and over 150 downstream genes^17^ Of primary concern for clinical gene editing is that off-target activity may result in cells with a proliferative advantage, increasing the potential for malignant transformation—a risk observed in early and recent gene therapy studies with viral vectors^18–20^.

Individual genomes differ from the human reference genome at approximately 4-5 million positions^21^. Furthermore, intraindividual somatic variation also occurs, although the frequency of these spontaneous variants is generally orders of magnitude lower than that of germline variants present in every cell within that individual^22^. Genetic variations such as single-nucleotide variants (SNVs), insertions, and deletions can overlap with Cas9 target-complementary or protospacer adjacent motif (PAM) sites. Such overlap can alter activity at existing off-target sites or create new ones. Thus, for exa-cel and similar therapies, a critical question is how genetic variation within patient populations may influence the risk of unwanted off-target editing.

As genome editing therapies expand to broader patient groups and different targets, accounting for human genetic diversity will be essential to ensure their consistent safety and efficacy across populations. Despite intense interest, the frequency and extent to which genetic variation generally affects cellular genome editing outcomes remain largely unknown. This major gap in knowledge is due to the lack of scalable, unbiased methods to experimentally measure genome-wide on- and off-target editing activity in cells from large, diverse populations, as well as tools capable of comprehensively assessing sufficiently diverse off-target sites to acquire fundamental understanding. Although previous studies have shown that genetic variation can affect genome editing and off-target activity, they have primarily focused on *in silico* predictions of variant-containing off-target sites rather than unbiased, genome-wide experimental assessments. Misek et al. found that genetic variation can confound the analysis of CRISPR/Cas9 viability experiments to identify ancestry-associated genetic dependencies for cancer cell survival^23^. However, this study does not directly measure either on- or off-target mutation frequencies, but instead indirectly quantifies changes in gRNA abundance as a proxy for on-target editing activity. A variant (rs114518452-C) in the intron of *CPS1* was identified by Cancelierri et al. to cause off-target editing associated with the exa-cel target at that genomic location^8^. Recently, Yen et al. confirmed that low-frequency off-target editing in the presence of this variant was detectable in two SCD patients treated with exa-cel, underscoring the importance of variation-aware off-target analysis^24^.

Experimentally unbiased methods (those that do not depend on *a priori* pre-selection of candidate sites) to define the frequency and location of unintended genome-wide off-target activity^9,25–29^ fall broadly into two categories: cellular methods that are more direct, and biochemical approaches that can be more sensitive. GUIDE-seq^9,30^ is a broadly adopted cellular method to define genome-wide on- and off-target activity of genome-editing nucleases such as Cas9, which was instrumental in defining the specificity of exa-cel^31^. It is based on the principle of efficiently integrating a short double-stranded oligodeoxynucleotide (**dsODN**) tag into nuclease-induced double-stranded breaks (**DSBs**), followed by tag-specific amplification of surrounding genomic DNA. CHANGE-seq is a biochemical method for selective sequencing of editor-modified DNA that is more sensitive than GUIDE-seq^25^, but requires further validation in cells. Improved versions of these assays with greater scalability or broader off-target coverage would enable a better understanding of how genetic variation affects genome editing.

Although early studies demonstrated that SNVs could influence Cas9 activity^32^, they could not systematically discern how prevalent or impactful these variants are in a population-scale cellular context. Previously, we used CHANGE-seq to demonstrate that SNVs can significantly affect Cas9 biochemical *in vitro* cleavage *across* six donors^25^. Computational studies leveraging data from the 1000 Genomes Project and Human Genetic Diversity Project estimated the theoretical prevalence of variant effects using off-target scoring algorithms^6,7^. Cancellierri et al. developed CRISPRme, a variant-aware off-target search tool, to identify off-target sites for the *BCL11A* enhancer target in exa-cel, and validated a variant-specific off-target in human hematopoietic stem cells that resulted in allele-specific indels and pericentric inversions^8^. A common limitation of *in silico* off-target studies is their reliance on genomic databases predominantly composed of individuals of European ancestry, potentially underestimating off-target risks in other populations^33^.

Another critical barrier to accurate bioinformatic prediction of variant effects on genome editor activity has been the lack of large-scale, diverse experimental datasets for training accurate models. Limitations of current datasets include: 1) sparse representation of active off-targets within the human genomes, 2) low overall numbers of data points, and 3) uneven representation of off-targets due to constraints of human genome structure and function on the presence of homologous off-target sites.

To address the critical gap in understanding the impact of individual genetic variation on Cas9 cellular activity, we developed GUIDE-seq-2, a new, streamlined, high-throughput method for population-scale, genome-wide profiling of Cas9 off-target activity. We applied GUIDE-seq-2 to six gRNAs in lymphoblastoid cell lines from 95 individuals characterized by the 1000 Genomes Project^21,34^. We identified a high-confidence set of variants that affect off-target activity by multiplex targeted sequencing. Our results reveal that common genetic variants frequently overlap Cas9 off-target sites and alter cellular off-target activity.

To further investigate these effects, we developed a novel massively parallel biochemical assay, CHANCE-seq, that enabled deep combinatorial profiling and the identification of sequence contexts and features associated with off-target cleavage *in vitro*. Finally, we developed a deep neural network model, CHANCE-net, to accurately predict the effects of overlapping genetic variants on Cas9 cleavage activity, revealing frequent novel context-dependent synergistic effects (**Fig. 1**).

**Fig. 1.**
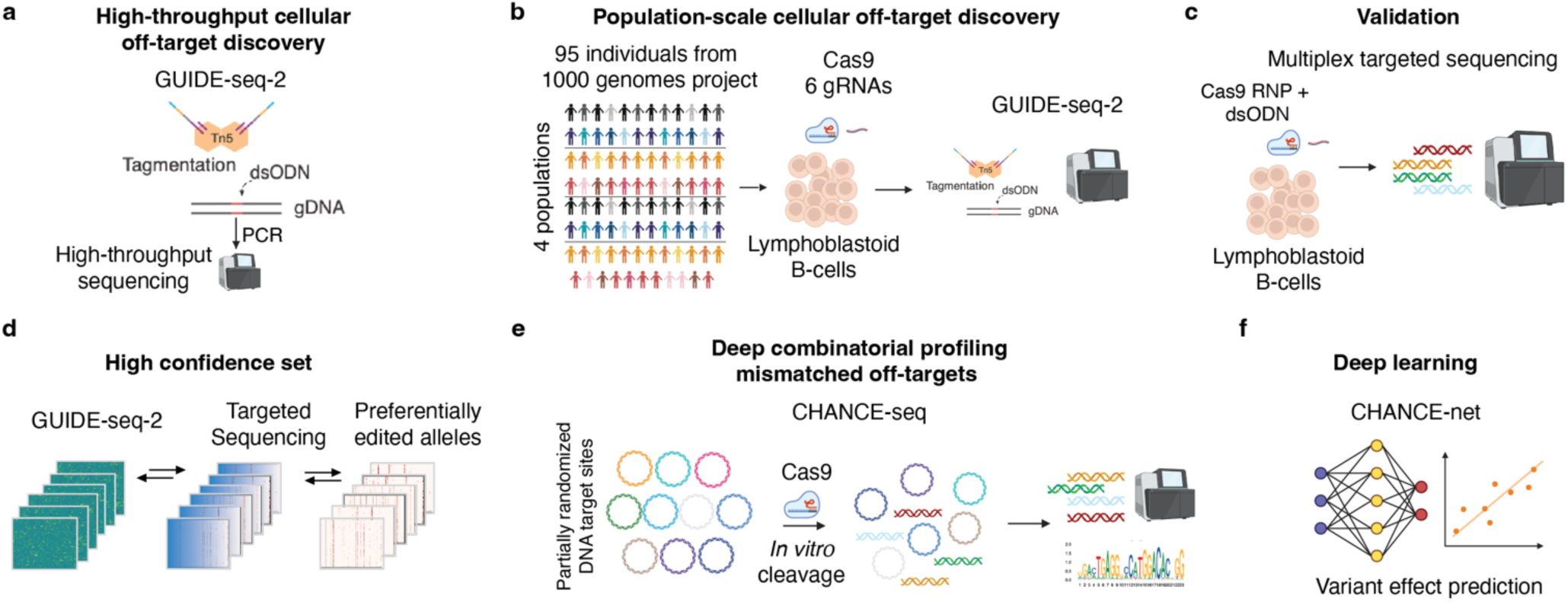
Understanding effects of human genetic variation on Cas9 genome-wide editing activity. Overview of the experimental design to quantify the frequency and magnitude of effects of individual genetic variation on CRISPR-Cas9 nuclease genome editing activity. **a**, First, we developed an optimized, streamlined, high-throughput GUIDE-seq-2 method to rapidly profile cellular on- and off-target editing activity. **b**, Second, we applied GUIDE-seq-2 on a population scale to discover Cas9 genome-wide editing activity associated with 6 gRNA targets in cells from 95 individuals from 4 populations characterized by the 1000 Genomes Project. **c**, Third, we performed multiplexed targeted sequencing to validate and quantify editing at the GUIDE-seq-2 sites identified in our population-scale survey. **d**, Fourth, by integrating orthogonal measures of variant effects, we defined a high confidence set of variant effects on Cas9 editing activity. **e**, Fifth, to more comprehensively survey the off-target landscape, we developed a massively parallel, combinatorial, biochemical assay, CHANCE-seq, to interrogate the biochemical activity of Cas9 across millions of mismatched off-target sequences. **f**, Finally, we trained a variant-aware, context-dependent machine learning model that accurately predicts the effects of genetic variation by distinguishing between neutral and synergistic mismatch effects.

## Results

### Development of a high-throughput method to evaluate cellular genome-wide activity of genome editors

To enable the analysis of Cas9 nuclease genome-wide off-target activity in a large number of well-characterized lymphoblastoid cell lines from genetically diverse individuals^34^, we reasoned that we could streamline GUIDE-seq^9,30^, as it would be challenging to scale the original method to many samples.

To substantially increase scalability, we searched for subtractive changes^35^ that yielded a streamlined, high-throughput method we called **GUIDE-seq-2,** which substantially reduces processing steps, time, and cost compared to the original method. GUIDE-seq-2 utilizes Tn5-transposon-based tagmentation^36–39^ and a new library design that eliminates the need for nested rounds of PCR (**Fig. 2a**), integrates community GUIDE-seq improvements^40^ without changing the original GUIDE-seq tag sequence to maintain full compatibility, and simplifies both library preparation and sequencing.

**Fig. 2.**
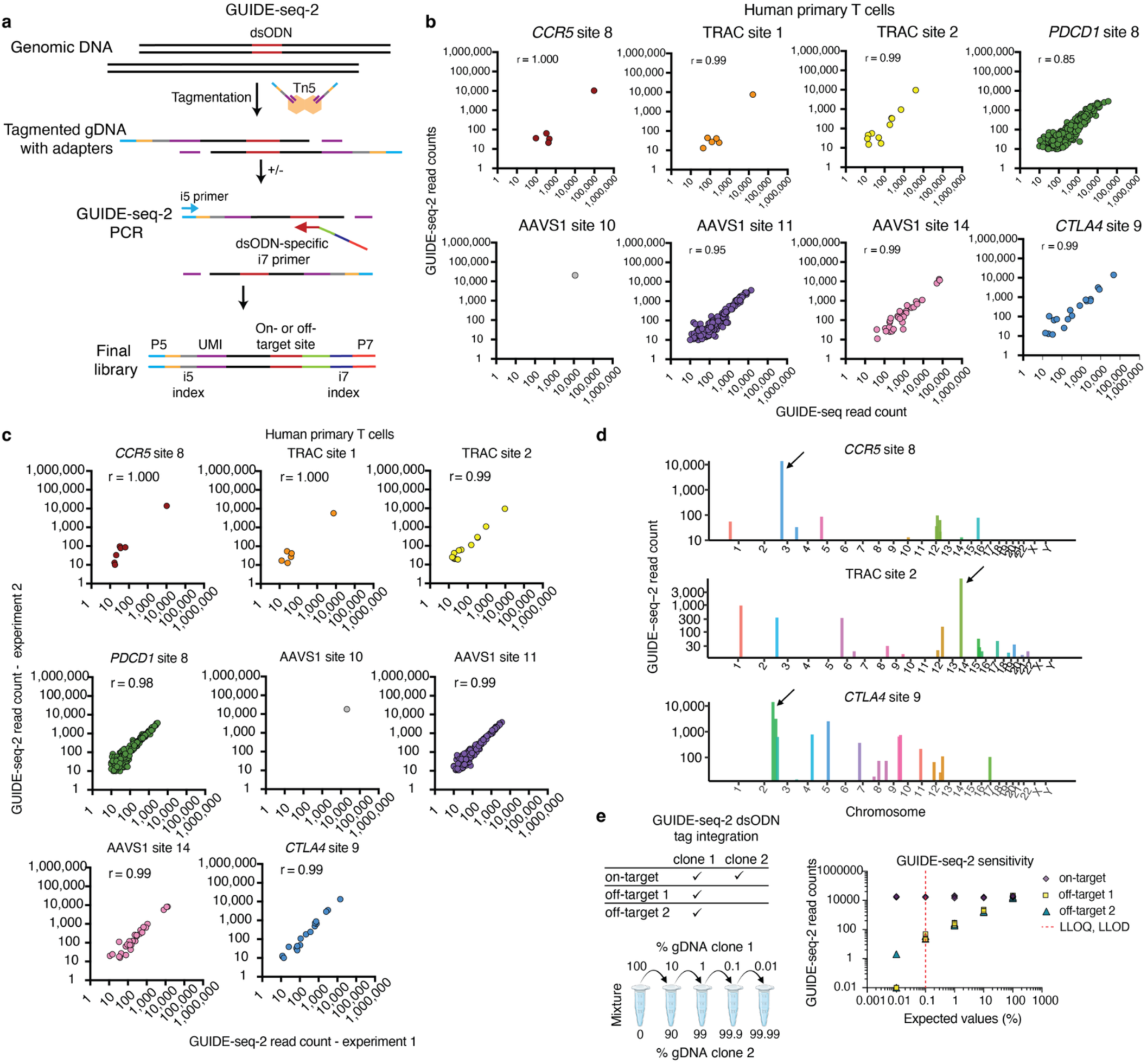
Development and optimization of GUIDE-seq-2, a highly scalable, sensitive, and unbiased method to define cellular genome-wide activity of genome editing nucleases. **a,** Schematic overview of streamlined GUIDE-seq-2 library preparation workflow. A short end-protected double-stranded oligodeoxynucleotide (**dsODN**) tag is integrated into sites of Cas9-induced DNA double-stranded breaks in living cells. Genomic DNA isolated from these cells is tagmented to provide a universal priming site. A single round of tag-specific PCR is performed to amplify tag-containing DNA for high-throughput sequencing. **b,** Scatterplots of GUIDE-seq and GUIDE-seq-2 read counts (log10) from experiments performed on primary human T cells for 8 target sites. **r** is Pearson’s correlation coefficient. **c,** Scatterplots of GUIDE-seq-2 read counts (log10) between two GUIDE-seq-2 libraries independently prepared from experiments performed in primary human T cells for 8 target sites, **r** is Pearson’s correlation coefficient. **d,** Manhattan plots showing the genome-wide distribution of GUIDE-seq-2 detected off-target sites with chromosomal position (x-axis) and read counts (y-axis). Arrow indicates the intended on-target site. **e,** Sensitivity of GUIDE-seq-2 established by determining lower limit of detection (LLOD) and quantification (LLOQ) X-axis: expected fraction of off-target in sample. Y-axis: GUIDE-seq-2 readcount. n=3.

Our redesigned GUIDE-seq-2 PCR strategy simultaneously improves detection accuracy by enabling mispriming detection, and sequencing without custom primers and sequencer configuration that were previously required. By introducing tagmentation, which simultaneously fragments genomic DNA and adds required sequencing adapters, we eliminated multiple library preparation and purification steps that include random genomic DNA shearing, end repair/A-tailing and adapter ligation, and reduced genomic DNA input requirements by 4-fold (**Extended Data Fig. 1, Supplementary Note 1, and Supplementary Protocol 1**). All experiments for GUIDE-seq-2 development were performed in primary human T cells.

We performed GUIDE-seq-2 and GUIDE-seq at 8 Cas9 target sites across 5 therapeutically relevant *loci* in CD4^+^/CD8^+^ human primary T cells (*CCR5*, TRAC, *PDCD1*, AAVS1 and *CTLA4*) with variable numbers of known off-targets identified in our earlier study^25^. We found a strong correlation between GUIDE-seq-2 and GUIDE-seq read counts at these 8 sites (*r* = 0.8 to 1.0, *r* is Pearson’s correlation coefficient) (**Fig. 2b**). For each of the 8 targets, we performed independent GUIDE-seq-2 library preparations and found that replicate read counts were strongly correlated (*r* = 0.99 to 1.0) (**Fig. 2c**), demonstrating high technical reproducibility. The positions of on- and off-target sites were genome-wide as expected (**Fig 2d**). We further estimated GUIDE-seq-2 sensitivity using defined mixtures of genomic DNA from ‘on-target’ single-cell clones with only on-target GUIDE-seq-2 tag integrations with DNA from ‘off-target’ clones with both on- and off-target tag integrations. We found that the lower limit of detection (LLOD) was between 0.01 and 0.1% of dsODN integrated cells (n=3). The lower limit of quantification (LLOQ) was approximately 0.1% of dsODN integrated cells (**Fig 2e**).

To filter out mispriming artifacts, in the GUIDE-seq-2 design we kept the original tag sequence but reduced the length of the primer annealing to the dsODN tag region, allowing us to observe expected sequences of 5 to 12 bases of dsODN region beyond the primer, and properly identify the expected junction between dsODN and flanking genomic DNA in the sequencing data (**Extended Data Fig. 1b**). An advantage of shortening the tag-specific primer (instead of increasing the tag length^40^), is that tag integration rates upon nucleofection remain unchanged, in contrast to the decreases in tag integration and sensitivity that can occur when tag length is increased^41^.

To further streamline the GUIDE-seq-2 workflow, we introduced a bead-based normalization step, using limiting amount of beads for single-step PCR purification and DNA normalization^42^, avoiding a labor-intensive serial dilution step for library quantification. Additionally, we adapted and optimized GUIDE-seq-2 for an automated liquid handling platform, demonstrating high technical reproducibility and comparability to manual GUIDE-seq-2, and expanding its potential for high-throughput applications (**Extended Data Fig. 2a-e**). In summary, our GUIDE-seq-2 approach is comparable to the original GUIDE-seq in sensitivity, is highly reproducible and automatable, requires less genomic DNA for library preparation, reduces processing steps and time to 3 hours, and has improved detection accuracy (**Supplementary Protocol 1**).

### CRISPR-Cas9 cellular off-target editing sites frequently overlap human genetic variants

GUIDE-seq-2 enables efficient and precise assessment of cellular Cas9 off-target activity, allowing investigation of the impact of human genetic variation at an unprecedented scale. Specifically, we analyzed cellular on- and off-target sites associated with six gRNA targets and one control from 95 donors across four populations, in a total of 665 GUIDE-seq-2 libraries (**ASW**, African Ancestry in Southwest USA; **CEU**, Utah residents with Northern and Western European ancestry; **CHB**, Han Chinese in Beijing; and **MXL**, Mexican Ancestry in Los Angeles, **Supplementary Table 2**, cell lines^21^). The six gRNA target sites (AAVS1 site 14, *CCR5* site 8, *CTLA4* site 9*, CXCR4* site 8, *LAG3* site 9, and TRAC site 1) were selected for this study due to their intermediate levels of off-target activity as measured in our previous study^25^ (**Supplementary Table 1**, gRNAs). GUIDE-seq-2 dsODN integration was optimized for lymphoblastoid cell lines (**LCLs**) characterized by the Genome-in-a-bottle consortium (**GIAB**)^43^ and evaluated at two targets with GUIDE-seq-2 (**Extended Data Fig. 2 f-j**). As a control, to identify the background of spontaneous, naturally occurring DNA DSBs, cells from each donor were transfected with dsODN tag only (without Cas9 or gRNA).

We identified 1,115 unique on- and off-target sites genome-wide, ranging from 56 to 333 per target (**Fig. 3a**), using stringent criteria that require at least two molecularly distinct GUIDE-seq-2 dsODN tag integration events for off-target site calling. These sites were distributed mostly in intergenic and intronic areas that comprise the majority of the human genome, and were also found in promoter, 5’ and 3’ UTR, and exonic regions in expected proportions (**Fig. 3b**).

**Fig. 3.**
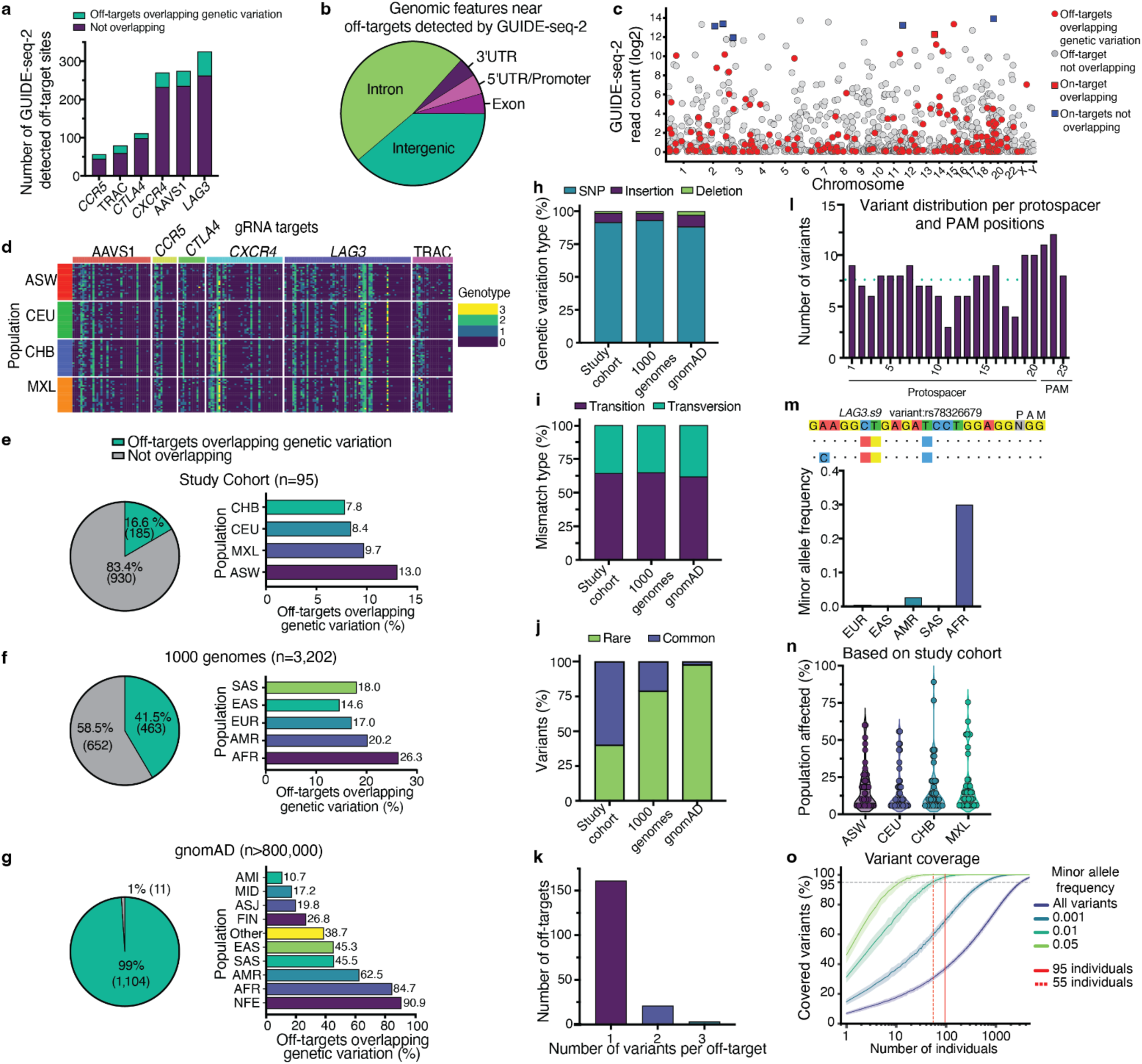
Cellular off-target editing sites detected by GUIDE-seq-2 frequently overlap genetic variants. **a,** Stacked bar plot of number of GUIDE-seq-2 on- or off-targets sites for each of 6 gRNA targets and proportion of sites that overlap with genetic variation, n=95 individuals. **b,** Pie chart of genomic distribution of off-targets detected in our study cohort. Untranslated Region (UTR). **c,** Manhattan plot showing off-targets with chromosomal position versus GUIDE-seq-2 read count, highlighted by presence of genetic variation. **d,** Heatmap of genotypes of variant-containing off-targets found in our study cohort. Horizontal rows represent individuals, Vertical columns represent one on- or off-target site. Color represents off-target genotype: off-target of homozygous reference (0), heterozygous (1), homozygous alternative (2), multiple variants with multiple combinations (3). African Ancestry in Southwest USA (**ASW**), Northern Europeans from Utah (**CEU**), Han Chinese in Beijing, China (**CHB**), Mexican Ancestry in Los Angeles, CA (**MXL**). **e,** Pie chart representing fraction of cellular off-target sites that overlap genetic variants and bar plot showing number of genetic variants per population in this study cohort (n=95 individuals) **f,** Same analysis as e, extending to all variants in the 1,000 Genomes Project (n=3,202). South Asian (**SAS**), East Asian (EAS), (**EUR**), Admixed American (**AMR**), African/African American (**AFR**) and **g,** Same analysis as e, extending to all variants in gnomAD (n>80,000), Amish (**AMI**), Middle Eastern (**MID**), Ashkenazi Jewish (**ASJ**), Finnish (**FIN**), Non-Finnish European (**NFE**) **h,** Bar plot indicating the proportion of genetic variants that contained deletions, insertions or SNPs in this study (n=212 overlapping variants), 1,000 Genomes Project (n=622), and gnomAD database (n=6,311) **i,** Bar plot indicating the proportion of SNPs that were transitions or transversions in each group **j,** Bar plot indicating the proportion of genetic variants that were common or rare in each group. rare=minor allele frequency (**MAF**) less than 1%. **k,** Bar plot showing on-/off-target sites overlapping with different numbers of variants **l,** Bar plot showing distribution of variants across the protospacer and PAM sequence. Turquoise dotted line represents average number of variants across all positions. **m,** Bar plot showing an example of a variant found predominantly within one genetic superpopulation. Visualization showing the on-target, reference variant and non-reference variant at position 2. **n,** Violin plot showing an estimated percentage of people affected by common variant-containing off-targets. **o,** Line plot showing percentage of variants covered based on bootstrap sampling from the 1000 Genomes Project (n=3202 individuals). Color represents different variant set: The gray dotted line represents 95% variant coverage. The red solid line marks our study size (n = 95). The red dotted line marks the sample size required to achieve 95% variant coverage for common variants. X-axis represents number of individuals drawn on log10 scale.

Cellular off-targets detected by GUIDE-seq-2 frequently overlapped human genetic variants (**Fig. 3c-d, Extended Data Fig. 3, Supplementary Table 3)**. Specifically, cumulatively 16.6% of on- or off-target sites (185 of 1,115) overlapped with one or more genetic variants found in our cohort (n=212 overlapping variants), with individuals of ASW ancestry having the highest frequencies of off-targets overlapping genetic variation (13%), and those of CHB ancestry having the lowest (7.8%) (**Fig. 3e**). When considering the entire 1000 Genomes Project population of 3,202 individuals with high-coverage whole genome sequencing (n=622 overlapping variants), the overlap percentage increased to 41.5% (463 of 1,115 sites) (**Fig. 3f**). Finally, when considering all variants in gnomAD (Genome Aggregation Database, n=6,311 overlapping variants), currently the largest collection of human genetic variation data^44^, we found that nearly all (i.e., 99% or 1,104 of 1,115) of sites overlap with at least one annotated genetic variant (**Fig. 3g**).

Among these variants, SNPs are the most frequent type, followed by insertions then deletions (**Fig. 3h**). Among the SNPs, transition mutations outnumber transversion mutations by an approximately a 2:1 ratio, consistent with expectations for genome-wide studies^45^ (**Fig. 3i**). Overlapping variants have a wide range of frequencies across populations. In our cohort, 60% variants are common variants (defined as minor allele frequency or MAF≥1%) (**Fig. 3j**). As expected, more rare overlapping variants were found as variant database size increased, with 79% rare variants in the 1000 Genomes Project and 98% in gnomAD.

Most GUIDE-seq-2 detected editing sites (n=161) overlapped with only one variant, although 24 off-target sites harbored two or three (**Fig. 3k)**. We observed that overlapping SNPs were evenly distributed across the protospacer and PAM positions (**Fig. 3l**) with an average of 7.6 SNPs per position. Interestingly, some variants were highly population-specific, suggesting the importance of population-specific, variant-aware analyses. For example, SNP rs78326679, occurring at the second position in *LAG3* site 9 off-target, is much more likely to occur in people of African American ancestry (**Fig. 3m**). We estimated that the average common off-target affecting variant will be found in 16.2% of individuals, however, this number will depend on the variant allele frequencies (**VAF**) observed in each specific population (**Fig. 3n**). For a more comprehensive estimate, we expanded our analysis to 58 targets and 1,403 off-targets analyzed by GUIDE-seq^25^, then looked for overlap of common variants in the 1000 Genomes Project. The average, common, off-target affecting variant was found in 22.1% of individuals (**Supplemental Fig. 1a**). When stratifying by targets with high (>100), medium (20 to 100), and low (<20) numbers of off-targets, no significant differences were found between groups **(Supplemental Fig. 1b)**. On a per individual basis, we estimate each person will have an average of 3% off-targets overlapping with their own common variants. **(Supplemental Fig. 1c)**.

Overall, our study of 95 individuals efficiently surveyed 97% of common variants and 34.1% of all variants in the 1000 Genomes Project (with MAF down to 0.03%) that overlapped the GUIDE-seq-2 cellular off-target sites discovered in this study (**Fig. 3o**). Based on our results, we estimate that on average, a sample of 55 individuals would be sufficient to survey 95% of common variants that overlap cellular off-target sites (**Fig. 3o**).

### Validation of GUIDE-seq-2 discovered off-target sites

To validate GUIDE-seq-2 discovered off-target sites and identify a high-confidence set of variant effects on off-target editing activity, we conducted multiplexed targeted sequencing to measure editing frequencies at 283 on- and off-target sites associated with the 6 targets we studied. This panel was designed to (1) cover a broad range of editing activities for regions without genetic variation based on GUIDE-seq-2 read counts; and (2) include all sites containing genetic variation. In this panel, 149 sites overlapped with 169 variants, while the remaining 134 sites were located in regions without genetic variation. Unedited controls were included for each site and each donor **(Extended data Fig 6).** Then, we assessed the significance of variant effects on editing activity as measured by indel and GUIDE-seq-2 tag integration frequency (**Fig. 4a**).

**Fig. 4.**
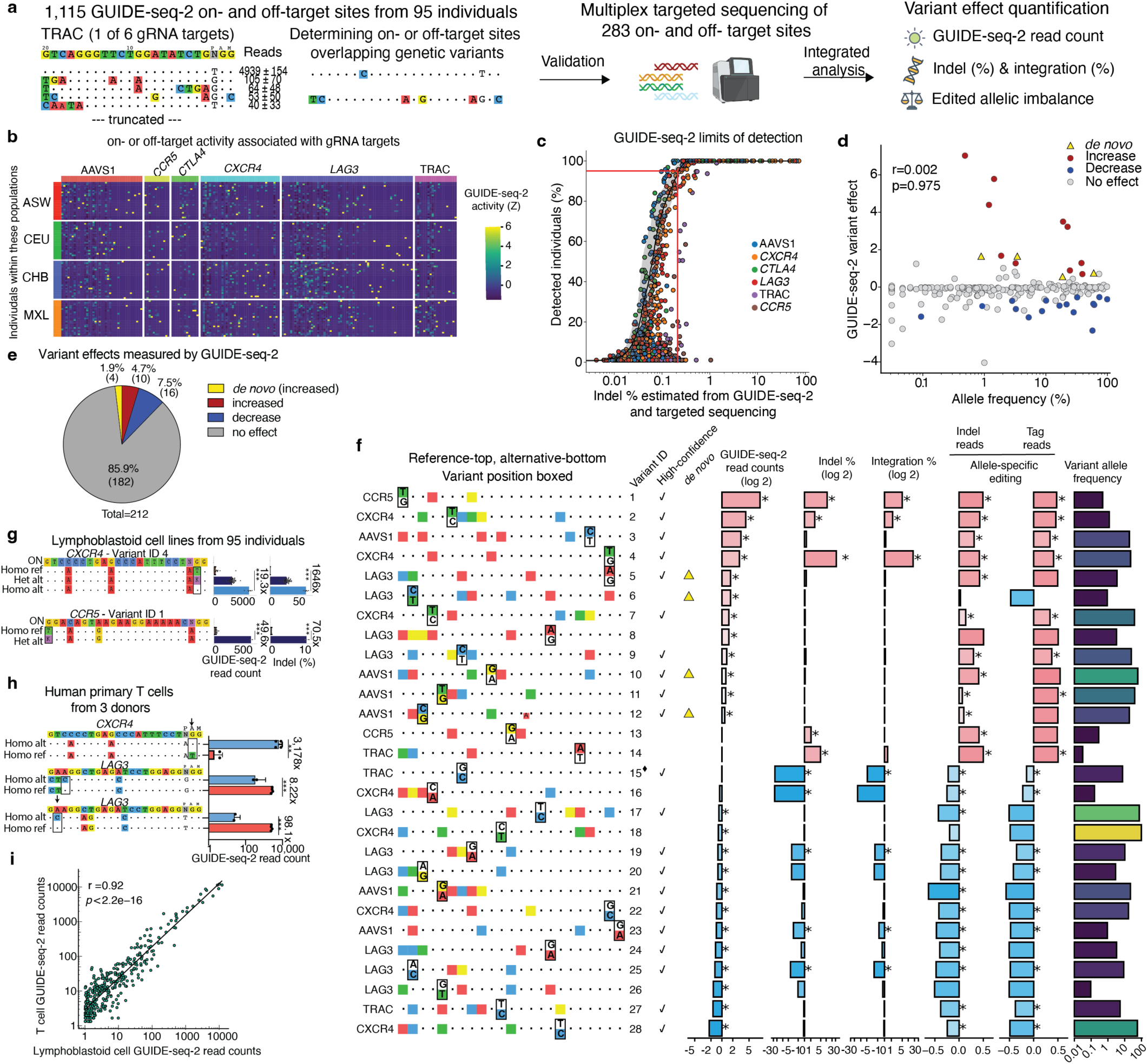
Overlapping genetic variants frequently affect editing activity. **a**, Schematic of validation experiments. On- and off-targets sites were identified by GUIDE-seq-2, a subset were validated using multiplex targeted sequencing. Statistical analyses were performed to quantify variant effect across orthogonal measures. **b**, Heatmap of GUIDE-seq-2 activity data, colored by Z-score, for 6 gRNA targets in 95 individuals across four 1000 Genomes Project populations: ASW, CEU, CHB, and MXL. Each row represents an individual, and each column represents an on- or off-target site. **c,** Scatterplot showing GUIDE-seq-2 limits of detection across 95 individuals. Each dot represents an on/off-target site. y-axis represents percentage of individuals having the on-/off-target. x-axis represents normalized GUIDE-seq-2 activity based on on-target indel frequency. Red line represents the GUIDE-seq-2 limits of detection for off-target sites (without genetic variants) detected in 95% of individuals with ∼0.2% indel frequency. **d**, Scatter plot showing allele frequency of the 212 variants detected by GUIDE-seq-2, colored by variants that are *de novo*, increase, or decrease editing activity r=Pearson’s. **e,** Pie chart showing proportion of 212 variants detected by GUIDE-seq-2 that are *de novo*, increase, or decrease editing activity. **f,** A detailed view of variants, defined by GUIDE-seq-2 and multiplexed targeted sequencing data, sorted based on GUIDE-seq-2 read counts. Off-target sequence alignment is shown to the right of the on-target gRNA names. Colored boxes represent mismatches to on-target sequence. Variant positions are boxed in black with alternative (non-reference) allele on the bottom. *De novo* and high-confidence variants are shown with a check mark. Variant 12 has a 1-bp insertion, indicated by a smaller red square. ♦On-target sequence contains a variant. Bar plots show variant effects in the heterozygous individuals. n=95 for GUIDE-seq-2, indel, integration analyses. Sample size varies based on the number of heterozygous individuals for variant allele imbalance analyses. Red represents increased effect and blue represents decreased effect. **g,** Detailed bar plots of unexpected and expected variant effect examples. On-target site shown above, mismatched nucleotides are colored. Variant nucleotide boxed. Student’s t-test and fold change are calculated between homozygous reference and homozygous alternative. ***p ≤0.0001. **h,** Bar plot of variant effects in human primary T cells. Fold change between homozygous reference (red) and homozygous alternative (blue) are shown to the right. **p ≤ 0.001 ***p ≤ 0.0001 **i,** correlation between human primary T cells, (3 donors, 2 replicates each) and LCLs, 95 donors, for off-targets without variants, r= Pearson correlation coefficient.

Using GUIDE-seq-2, we quantified a broad range of off-target activities associated with 6 gRNAs targets that varied substantially between individuals from the ASW, CEU, CHB, and MXL populations (**Fig. 4b**). We confirmed that GUIDE-seq-2 read counts were strongly correlated with indel and GUIDE-seq-2 tag integration frequencies as measured by multiplex targeted sequencing (**Extended Data Fig. 4a**, **Supplementary Table 4**). We estimated GUIDE-seq-2 lower limits of detection in our studies to be ∼0.2%, based on the 930 sites not overlapping with any variants in the 95 donors and the lowest estimated indel frequencies of off-target sites we could detect in 95% of samples (estimated by the product of GUIDE-seq-2 relative off-target activity and on-target editing percentage) **(Fig. 4c)**.

### Overlapping genetic variants frequently affect off-target editing in human cells

To assess variant effects on off-target activity, we integrated five variant effect measures. Specifically, we modeled the effect of overlapping genetic variation on normalized GUIDE-seq-2 read counts, indel frequencies, tag integration frequencies, and allele-specific editing measured as indels or tag integrations. If Cas9 exhibits a preference for reference or alternative genotypes, this should be reflected as an imbalance in the frequency of edited alleles (Extended data **Fig. 5a**). We defined a high-confidence variant effect as one that was significant in two or more of these measures (see **Methods**).

We found no significant correlation between variant allele frequency and GUIDE-seq-2 variant effect (r=0.002, p-value=0.9) **(Fig. 4d).** A high proportion of variants (14.1%) measured by GUIDE-seq-2 significantly impacted off-target activity, with at least two-fold read count change, of which 6.6% (n=14) and 7.5% (n=16) of variants led to increased or decreased editing, respectively. Of the 14 variants that increased editing, 4 were *de novo* sites in which variants created new off-target sites not detected in the reference genome **(Fig. 4e)**. Variant effects on GUIDE-seq-2 reads ranged from –2.3 to 7.0 (log2-transformed GUIDE-seq-2 read counts) and were corroborated by comparable changes in indel and tag-integration percentages (**Fig. 4f, Extended Data Fig. 5b, Supplementary Table 3**). Notably, variant effects on TRAC site 1 on-target activity (high-confidence variant #15) and variants #13, 14, and 16 were only identified by targeted amplicon sequencing, as it was at the upper limits of quantification by GUIDE-seq-2 **(Fig 4f, Extended Data Fig 6).** Additional variants that had significant GUIDE-seq-2 read counts but no targeted sequencing data available, as well as those variants with deletions can be seen in **Extended data Fig 5b and 6**.

Certain variant effects were unexpected, such as a T>G variant at the first position of a Cas9 off-target sequence associated with a *CCR5*-targeting gRNA (high-confidence variant #1), which surprisingly led to a substantial 70-fold increase in Cas9 indels to ∼10% (**Fig 4f-g, Extended Data Fig. 6**). Our findings underscore the complexity of predicting Cas9 activity on variant-containing genomic sites and highlight the value of high-throughput profiling techniques.

Others were more predictable, such as those that revert a mismatch to the canonical *S. pyogenes* Cas9 NGG PAM sequence. For example, a T>G variant that altered an NGG PAM-containing off-target sequence (associated with a gRNA targeted to *CXCR4,* high-confidence variant #4) to NTG significantly reduced Cas9 editing activity, consistent with the importance of PAM recognition to license Cas9 target DNA cleavage^46^ (**Fig 4f-g, Extended Data Fig. 6**).

### Strong variant effects on off-target editing observed in human primary T cells

To determine whether our results extended to therapeutically relevant primary human cells, we performed GUIDE-seq-2 in primary human T cells from three healthy donors and evaluated Cas9 off-target activity for two gRNAs target sites (*CXCR4* site 8 and *LAG3* site 9). By directly evaluating GUIDE-seq-2 read counts correlations between the three donors (**Extended Data Fig. 4b**), we found strong evidence that the genome-wide off-target activity of Cas9 in these human primary T cells is affected by the presence of SNVs (**Fig. 4h**) with approximately 8 to 3,000-fold changes in GUIDE-seq-2 read counts. To understand how cell-type influenced editing outcomes, at off-target sites that did not overlap with genetic variants, we compared GUIDE-seq-2 on- and off-target read counts from our 95 LCL donors and 3 T-cell donors for 3 target sites (*CXCR4* site 8, *LAG3* site 9, and *CCR5* site 8). GUIDE-seq-2 read counts were strongly correlated between LCLs and T cells (Pearson correlation coefficient of 0.92), suggesting that our data generated in LCLs are generalizable to other cell types **(Fig. 4i).**

### Creation and validation of a massively parallel biochemical assay to quantify activity at diverse mismatched off-targets

Our population-scale GUIDE-seq-2 analysis identified, to our knowledge, the largest set of genetic variants known to affect Cas9 genome-wide off-target activity and revealed the first estimates of the frequency with which overlapping off-targets modulate Cas9 activity. However, it would be challenging to infer general characteristics of sequence contexts and genetic variants likely to impact Cas9 activity based on 28 variant-affected off-target sites alone.

To better understand influential genetic context and mismatches, we designed a high-throughput, cost-effective, biochemical assay that could measure Cas9 activity within highly diverse, partially randomized, off-target libraries which we called Combinatorial High-throughput Analysis of Nuclease Cleavage Efficiency by Sequencing or **CHANCE-seq** (**Fig. 5a, Extended Data Fig. 7a**). We reasoned that quantitative measurements of Cas9 activity at millions of mismatched target sequences could be more informative than earlier ensemble measurements of larger libraries^47^, where library members are statistically unlikely to be shared between control and treatment conditions and, therefore, activity quantification at individual targets is not possible. These diverse quantitative measurements could subsequently be used to train deep learning models to predict the impact of any genetic variant on Cas9 activity.

**Fig. 5.**
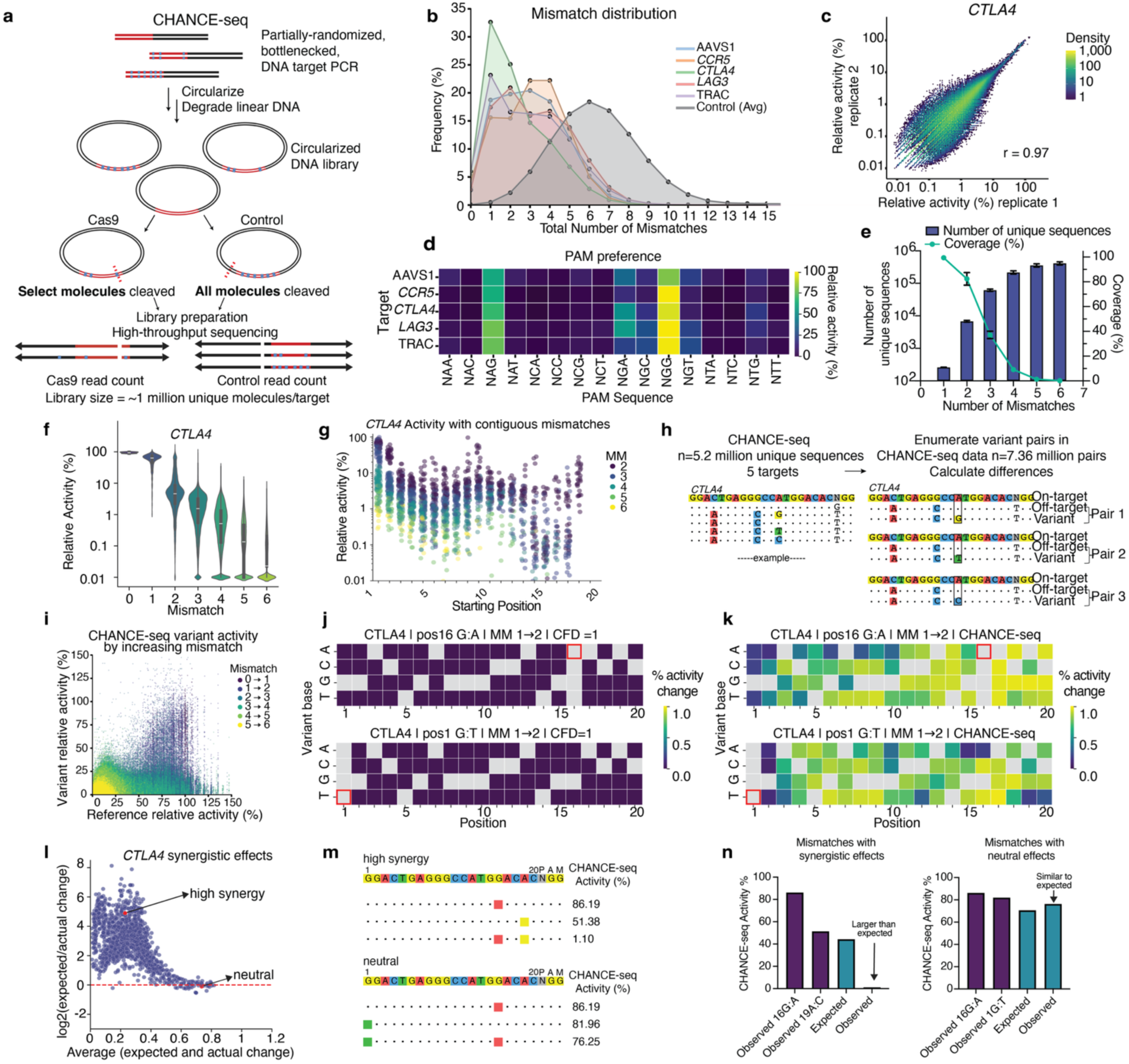
Deep combinatorial profiling of millions of mismatched targets with CHANCE-seq reveals frequent, context-dependent, synergistic variant effects on Cas9 activity. **a,** Schematic of CHANCE-seq assay which creates libraries of partially randomized target sites and selectively sequences those that are cut by Cas9 compared to a control. Red indicates target, black is plasmid backbone, blue squares are randomized mismatches in protospacer and PAM. The same partially randomized target sequences are analyzed across control and Cas9 treatments. A control restriction enzyme cleaves a site adjacent to all randomized targets, whereas only select molecules are cleaved with Cas9. **b,** Histogram of frequency of mismatches in CHANCE-seq libraries treated with Cas9 versus control libraries. Average of two Cas9 technical replicates for each target and five control libraries **c,** Binned scatterplot of % relative activity between replicate 1 and replicate 2 for Cas9 CHANCE-seq library for *CTLA4*, colored by density. *r* = Pearson’s. **d,** Heatmap of PAM specificity from inferred from CHANCE-seq data, colored by percent relative activity. **e,** Bar plot showing number of unique sequences at each mismatch level in CHANCE-seq, the line shows percent coverage of unique sequences. Error bars are standard deviation of 5 targets in control libraries. **f,** Violin plots showing relative activity % at each mismatch level 0-6 for *CTLA4* target using CHANCE-seq. **g,** Strip plot illustrating contiguous mismatch start positions (2-6 mismatches) with % relative activity on a log scale. Only longest contiguous block (length of block equal to number of mismatches) is shown. **h,** Schematic overview of CHANCE-seq variant pairs analyzed. **i,** Scatterplot of effect of single mismatches on relative activity in variant pairs, colored by mismatch. **j-k,** Heatmaps show differences in activity when increasing the number of mismatches from 1 to 2 for CFD (**j**) and CHANCE-seq (**k**). Both reference and variant sequences have one prior mismatch at each of 19 other positions, but variant sequences have one additional variant mismatch at pos16:G→A (top) or pos1:G→A (bottom) (highlighted in red). The percent activity change is calculated as (ref-var)/ref. **l**, MA plot illustrating synergistic effects of combined mismatches. Each dot represents a pair of mismatches. The x-axis shows the mean of the expected and observed activity changes, and the y-axis shows the log2(expected/observed) ratio. A pseudocount of 0.001 was added to both expected and observed activity changes. Positive values indicate synergistic effects, where combined mismatches lead to a greater reduction in activity than expected under the independence assumption, while points near zero represent neutral effects. **m**, example *CTLA4* sequences of mismatches with high synergy verses those with neutral effects with activity values. Sequences chosen from **5l**. **n,** barplot with expected activity change vs observed when there is high synergy vs neutral mismatch effects.

CHANCE-seq builds on principles of CHANGE-seq, a method we previously developed to selectively sequence DNA cleaved and linearized by Cas9 from populations of circularized genomic DNA molecules^25^, but operates on large, partially randomized off-target libraries instead of genomic DNA. CHANCE-seq enables profiling of Cas9 activity at millions of mismatched targets with many higher-order combinations of 3 or more mismatches. CHANCE-seq libraries can be orders of magnitude larger than earlier oligonucleotide array-based approaches^48,49^, overcoming limitations that off-target sites identified from genomic DNA are usually small in number, therefore have sparse representation of active off-targets within the human genome, and do not contain sufficient mismatch combinations to train machine learning models.

Using CHANCE-seq, we surveyed five of the targets (AAVS1, *CCR5*, *CTLA4*, *LAG3*, and TRAC) that we profiled with population-scale GUIDE-seq-2. First, we generated highly diverse libraries (10^10^ or more) where each position of a 23 bp Cas9 target and PAM sequence has been partially randomized by PCR, resulting in unique molecular index (**UMI**) barcoded libraries of off-targets with an average of 6 mismatches when compared to the original on-target sequence (**Supplementary Table 5**, **Fig. 5a**). The presence of UMIs can be used to correct for sequencing errors and other artifacts that could confound analyses. Second, critically, to generate circularized DNA libraries containing multiple copies of the same randomized target, we bottlenecked and amplified a subsample of approximately 1 million sequences. Third, we split and treated these circularized DNA libraries with either Cas9 that can recognize and cleave similar on- or off-target sites, or a control restriction enzyme that cuts a site adjacent to the randomized target sequence, cleaving all molecules. Only cleaved DNA circles have free ends for subsequent adapter ligation and high-throughput sequencing. For each on- or off-target sequence, we calculated the ratio of normalized CHANCE-seq read counts between Cas9 and control libraries as a measure of Cas9 activity (**Fig. 5a, Extended Data Fig. 7a**, **Supplementary Note 2, and Supplementary Protocol 2)**.

### Massively parallel CHANCE-seq measurements are globally consistent with Cas9 activity preferences

To determine whether our CHANCE-seq experiments effectively enriched for Cas9-cleaved DNA molecules, we compared the frequency of mismatches within the library of partially randomized target sequences when cleaved by Cas9 versus a control restriction enzyme that cuts all library molecules. We observed that Cas9-cleaved libraries were significantly enriched for sites with fewer mismatches compared to control libraries, 0.05, indicating that the assay was functioning as intended (**Fig 5b**).

Relative off-target activity measured between CHANCE-seq technical replicates was strongly correlated (*r* = 0.89 to 0.9) (**Fig. 5c, Extended Data Fig. 7b**). Cas9 PAM preferences determined by CHANCE-seq showed highest activity (94.9% ± 8.5%) at canonical NGG PAM, some activity at NAG (69.3% ± 7.8%) and NGA (35.5% ± 17.2%), and minimal activity at other non-canonical PAMs (**Fig. 5d**). The coverage of sequences with up to 6 mismatches ranged from 0.9 to 1.2 million sequences for each target, after filtering for adequate representation in control libraries (**Fig. 5e**). Off-target activity typically decreased with increased mismatch number, although we observed a long tail of exceptions where high mismatch sites retained high activity (**Fig. 5f, Extended Data Fig. 7c**).

The longest blocks of contiguous mismatches that preserved high activity were found in PAM-distal regions (starting at 5’ positions 1-2 as expected), but also more surprisingly in the middle (starting at positions 9-11) (**Fig. 5g**, **Extended Data Fig. 7d**). These two regions where Cas9 is most tolerant to contiguous off-target site mismatches coincide with those where structural studies have found protein:DNA phosphate backbone contacts (between target strand positions 1-2, 8-9, 10-13) that stabilize off-target site unwinding and recognition^50^.

Transition mutations such as G>A or T>C that result in wobble base pairings (rG:dT or rU:dG) were generally well tolerated and had the smallest effect on activity **(Extended Data Fig. 7e)**, specifically at positions 4, 16 and 22 **(Extended Data Fig. 7f),** whereas Hoogsteen and transversion mutations were most deleterious for Cas9 activity **(Extended Data Fig. 7e)** particularly at positions 1, 22, and 23 **(Extended Data Fig. 7f).** In addition to the PAM, positions 16 and 18 appeared least tolerant to mismatches while maintaining activity. At position 16, there is a stark contrast between transition mismatches, which resulted in high editing activity and transversions that lowered activity (**Extended Data Fig. 8,9)**. At higher off-target mismatch numbers, high-activity sites often accumulate more PAM-distal mismatches and mismatches become less common in the seed region (**Extended Data. Fig. 9**). Many of the sequence patterns associated with low and high off-target activity are consistent with those reported in prior studies that analyzed low-mismatch off-targets harboring 1-3 mutations ^51^.

Due to the unique ability of CHANCE-seq to interrogate millions of off-targets, it enables quantitative assessment of variant effects in many diverse contexts by comparing activity differences between homologous off-targets. Specifically, we enumerated all pairs of off-targets with up to 6 mismatches that differed by a single mismatched nucleotide and defined single nucleotide variants (SNVs) as those that increase mismatches to their on-target sequence, resulting in 7.36 million variant target pairs (**Fig. 5h**). The majority of variants that increase the number of mismatches relative to the on-target site reduced Cas9 activity (**Fig. 5i**). Variants have a greater impact on activity (‘leverage’) when the total numbers of “off-target context” mismatches is low (e.g. 0-2) compared to those with higher mismatches (3-5) (**Fig. 5i**).

### CHANCE-seq reveals novel, context-dependent, synergistic effects of mismatches on Cas9 activity

CHANCE-seq enables the quantification of mismatch-associated activity changes in many different sequence contexts, with coverage of activity at mismatched targets increasing by more than 6,000-fold per target compared to earlier methods (**Fig. 5e**). The cutting frequency determination (**CFD**) score, a leading method for predicting off-target activity, was trained on gRNAs with a single-mismatch, insertion, or deletion, resulting in an average of 188 gRNAs per target. In contrast, CHANCE-seq experiments quantify on average 1 million mismatched sites per targetacross sequences with mismatch numbers ranging to 6 and beyond. CFD score relies on a penalty matrix that varies by position and mismatch identity, but assumes penalties are fully context-independent. The penalty for a mismatch at a given position does not change, regardless of whether there are neighboring mismatches. For example, for every type of mismatch at every protospacer position, CFD predicts that an additional position 16 G→A or position 1 G→T mismatch will have no impact on activity (**Fig. 5j**). In contrast, for any mismatch, CHANCE-seq activity values dynamically change based on position and mismatch identity of contextual mismatches (**Fig. 5k**). For the same mismatches that CFD predicts have no effect, we observed activity changes at many mismatch combinations (**Fig. 5k**). Quantifying mismatch activity in diverse sequence contexts enables identification of synergistic versus neutral mismatch combinations. Many mismatches, particularly those with modest effects, can exert synergistic effects when combined (**Fig. 5l**). For example, a mismatch can have strong synergistic effects with a nearby mismatch, but largely neutral effects with a distant mismatch (**Fig. 5m and 5n**). Our common observation of context-dependent effects of mismatches emphasizes the importance of large-scale, diverse training datasets produced by methods like CHANCE-seq for accurate machine learning model development.

### Creation and validation of CHANCE-net deep learning model to predict the effects of mismatches on Cas9 activity

Experimentally assessing over 90% of known genetic variants overlapping potential off-target sites for the six targets in this study would require analyzing cells from ∼2,000 individuals (**Fig. 3o**), which is impractical, even with high-throughput methods such as GUIDE-seq-2 for sensitive and unbiased off-target discovery. Inevitably, there will be variants that cannot be surveyed experimentally due to a lack of access to cellular donors with that genotype. Similarly, by design, our *in vitro* high-throughput assay, CHANCE-seq, surveys diverse partially randomized sequences rather than directly evaluating genomic off-targets. To bridge this gap, we developed a deep learning model to predict and characterize the full variant landscape associated with off-target activity.

By improving mismatched target coverage and developing an experimental approach capable of deeply quantifying context-dependent mismatch effects, CHANCE-seq addresses gaps in current off-target variant effect measurement and prediction methodologies. CFD predicts that certain mismatches at positions 14 and 16 will always completely eliminate Cas9 activity, and assigns a score of 0. While CHANCE-seq data confirms that this is largely true, it also identifies unexpected exceptions (**Fig 6a).** For example, CFD predicts total activity loss when a mismatch occurs at position 14 T>G (*CTLA4*), however, off-targets containing this mismatch still retained almost half of their activity in CHANCE-seq (**Fig 6b).** Similarly, there are 14 mismatches across positions 1, 5, 6, 7, 8, 11, and 16 that CFD predicts never have an effect and assigns a score of 1 regardless of contextual mismatches. Our CHANCE-seq data show that there are many sequence contexts in which variants at these positions have a wide range of effects (**Fig 6c).** In some cases, activity can be reduced by more than 20-fold (**Fig. 6d)**. These discrepancies likely arise because many current prediction methods, like CFD, are trained on single-mismatch data or on genomic DNA where off-target sites are sparsely represented. We reasoned that expanding the off-target dataset could mitigate blind spots observed in previous approaches and could lead to the development of a more accurate deep learning model for prediction of the impact of variants on Cas9 activity.

**Fig. 6.**
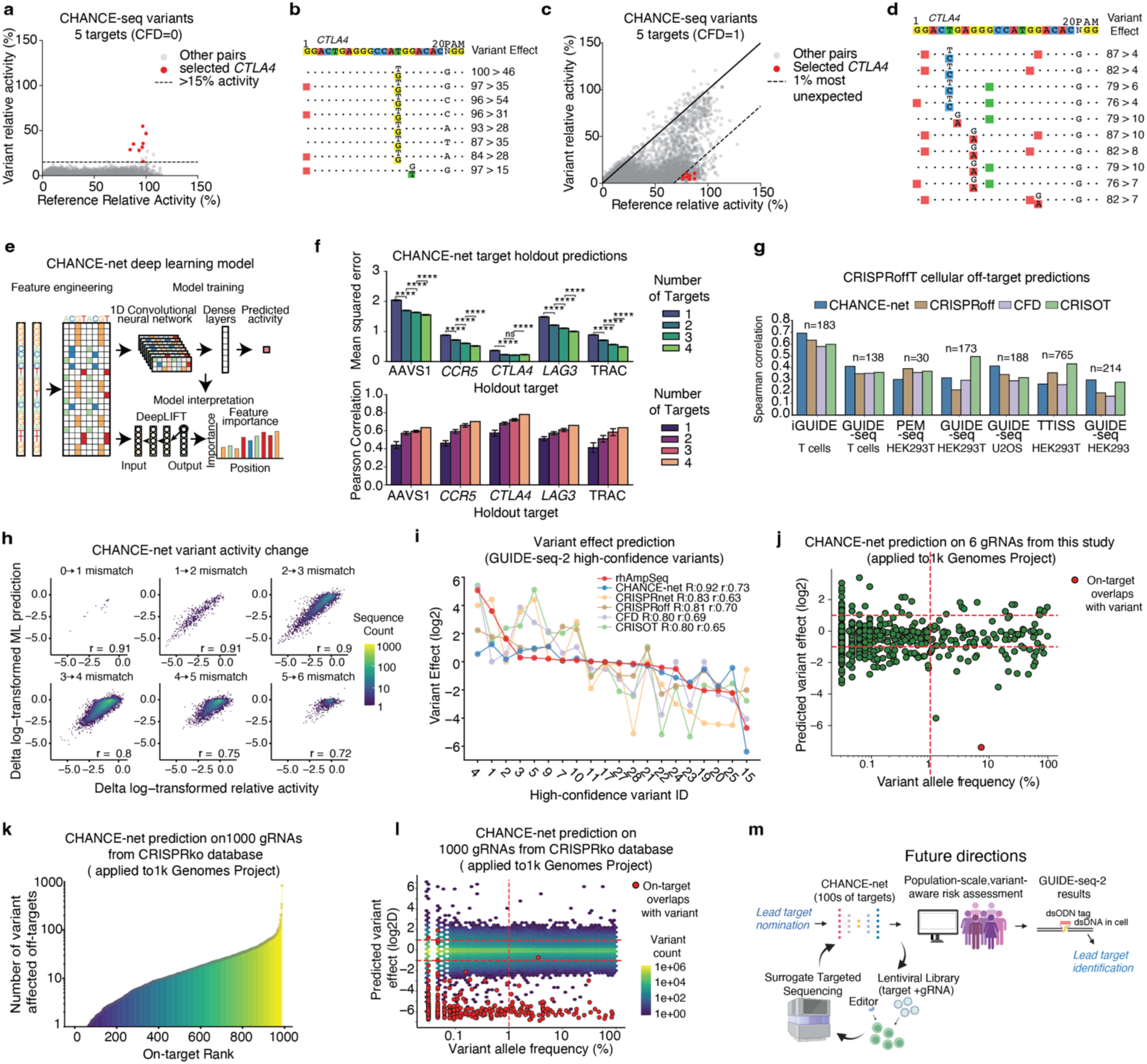
CHANCE-net convolutional neural network model enables accurate prediction of genetic variation impact in any sequence context. **a-c,** Scatterplot of reference and variant CHANCE-seq relative activity where CFD predicts either complete abrogation of activity (**a**) or no change in activity (**c**). **b,d,** Examples of unexpected observations for CTLA4 are highlighted in red and their sequences are shown with their percent relative activity. **e,** Schematic of deep learning framework. Input of 23-bp sequences from gRNA targets and off-target sites, encoded via one-hot encoding. They are processed by 1-D convolutional layers and dense layers to predict off-target activity. DeepLIFT is used for feature importance analysis. **f,** Bar plot of CHANCE-net model prediction performance on a hold-out target when trained on 1-4 other targets. For mean squared error, each point corresponds to one off-target sequence in the hold-out target. For Pearson correlation, each point corresponds to one hold-out experiment. **** = p ≤ 1.0 × 10⁻⁴; error bars = standard error. **g,** Bar plot benchmarking performance of CHANCE-net and 3 other predictive models on CRISPRoffT cellular off-target dataset. Spearman correlation is between predicted off-target activity and experimentally measured off-target activity across 7 cell types/assay groups. n = number of test off-target sequences in each group. **h**, Scatterplots of CHANCE-net prediction performance for variant activity. Subplots correspond to pairs of off-target sites that differ by a single mismatch, grouped by the number of mismatches. X-axis = CHANCE-seq delta log-transformed relative activity, y-axis = model predictions. r= Pearson. **i,** Line plot showing variant effect comparisons between rhAmp-seq, CHANCE-net, and four state-of-the-art CRISPR-Cas9 models. X-axis shows high-confidence variant ID referenced in Fig. 4f. R = Spearman and r = Pearson. **j,** Scatterplot of CHANCE-net predicted variant effects and the variant allele frequency based on 6 gRNAs used in this study. CHANCE-net Model applied to 1,000 genomes database off-targets with 23bp in length and SNP variants, red is on-target that overlaps with variant (n=516). **k,** dot and stem plot showing **988** gRNAs (Brunello library) on the y-axis and the number of variant affected off-targets per guide on the y-axis **l,** Scatterplot of CHANCE-net predicted variant effects and the variant allele frequency based on **988** gRNAs from **Brunello library**. (n=9.5 million. **m,** Schematic depicting vision for future utilization of GUIDE-seq-2, CHANCE-seq, and CHANCE-net for variant-aware risk assessment, enabling better lead target selection.

To comprehensively model off-target sequences and their activity, we trained a single convolutional neural network (CNN) model (called CHANCE-net) (**Fig. 6e**) to predict Cas9 activity, using 90% of the CHANCE-seq data from five gRNA targets for training and 10% as a held-out test set. The goal of this model is to predict Cas9 activity at unseen off-target sites, such as off-targets overlapping genetic variants in patients. The model performed well, accurately predicting Cas9 activity on off-target sequences with up to six mismatches (*r* = 0.81 to 0.96) (**Extended Data Fig. 10a**). To verify the model’s predictions, we applied an interpretable machine learning technique, DeepLIFT^52^, to analyze the CNN’s learned parameters. This feature attribution analysis revealed that mismatches at positions 5 to 7, 16, and 18 of the target sequence exerted the greatest influence on predicted Cas9 activity (**Extended Data Fig. 10b**). These areas correspond to target regions that may allosterically activate Cas9 cleavage^53^ and the nucleotides adjacent to the cleavage site, respectively, which have been previously reported in the literature.

To determine whether our model could generalize to off-target sites associated with unseen targets, we further trained it by leaving out one or more targets. When implementing leave-one-target-out training, we observed that mean squared error significantly decreased, and Pearson correlation increased as more targets were included in the training set (**Fig 6f)**. Although we anticipate that adding more target sites will likely further enhance CHANCE-net’s performance, we applied the current model to predict off-target activity across four cellular off-target assays and four distinct cell types from the CRISPRoffT database. (**Fig. 6g, Extended Data Fig 10c, d**). Our results show that even trained on a limited number of targets, CHANCE-net achieved the highest Spearman correlation (**Fig. 6g**) and highest Pearson correlation (**Extended Data Fig 10c**) in more than half of the datasets compared to CRISPRoff, CFD, and CRISOT models and had comparable prediction performance in remaining datasets (**Fig. 6g, Extended Data Fig 10c, d, Supplementary table 7**). Since CRISPR-net was trained on GUIDE-seq data, it could introduce data leakage; therefore, its predictions were excluded from the main analysis. However, CRISPR-net predictions are provided in **Supplementary Table 7** for reference.

### CHANCE-net deep learning model predicts the impact of genetic variants

Next, we assessed the model’s ability to predict variant effects. Using variant pairs generated as described in **Fig 5h** and 90% of CHANCE-seq data from five gRNA targets for training and 10% as a held-out test set, the model was able to capture Cas9 activity changes induced by SNVs within these off-target sequences (**Fig. 6h**). The model’s consistent accuracy for all targets suggests the potential for developing general predictors of genome editor off-target activity.

We next benchmarked our variant-aware off-target activity predictions against leading *in silico* predictors, using the high-confidence variant effects we previously identified and quantified by targeted sequencing as a test set. CHANCE-net predictionspredicitons were strongly correlated with variant-associated activity differences (Spearman’s R = 0.92, Pearson’s *r* = 0.73), outperforming other state-of-the-art methods, including CRISPRnet^54^, CRISPRoff^55^, CFD^56^, and CRISOT^57^ (**Fig. 6i**).

We applied our model to predict the effects of all naturally occurring variants reported in the 1000 Genomes Project based on the 1,115 on- and off-target sites identified in this study, associated with the 6 targets we studied. We identified 43 variants (allele frequency >= 1%) that could substantially affect activity at 42 off-target sites (>=1% activity difference) (**Fig. 6j**, **Supplementary Table 6**).

To gain a more comprehensive understanding of variant-affected off-target sites, we applied CHANCE-net to 988 gRNAs targeting 254 pan-cancer genes in the Brunello library^56^ across individuals from the 1000 Genomes Project. We found that each gRNA, on average, had 21.5 variant-affected off-target sites (median = 12) **(Fig 6k).** Plots of variant allele frequency versus predicted variant effect enable identification of common genetic variants most likely to increase off-target activity **(Fig 6l).** This analysis demonstrates the potential utility of CHANCE-seq in identifying lead targets that are least susceptible to new variant-associated off-target activity, underscoring its relevance for future research and therapeutic applications.

We envision two possible workflows integrating GUIDE-seq-2, CHANCE-seq, and CHANCE-net. First, as we have utilized in this paper, experiments can be performed on gRNAs of interest, and the model can enable a variant-aware *in silico* search for these specific guides. In the future, we anticipate that performing CHANCE-seq across hundreds of target sites and calibrating with cellular data will improve model performance to the point where CHANCE-net can predict variant effects for any target site without additional experimental data. This more generalizable model could accelerate therapeutic genome editor development by streamlining lead target nomination and off-target validation validation **(Fig. 6m)**.

## Discussion

### Discussion and future directions

Our study provides the first experimental, population-scale view of the impact of genetic variation on editing activity, and suggests that Cas9 cellular off-targets frequently overlap with common genetic variants and that variant effects on off-target activity are highly context-dependent. In a therapeutic context, of greatest concern would be the subset of variants that increase off-target editing and safety risks to patients^58,59^ (although it should be noted that only some unintended off-target sites may have adverse biological consequences). For biological research, genetic variation between donor cell lines or primary cells may confound genome-editing outcomes. Our estimate of frequent variant-induced effects on editing suggests that they should be carefully accounted for in the early stages of therapeutic lead target discovery or basic research, in a population-specific manner. This is further supported by recent reports of genetic variants that caused allele-specific off-target activity were observed in a SCD clinical trial^28^.

GUIDE-seq-2 can simplify cellular off-target analysis of genome editing for many labs: it enables large-scale experimental design, as in our study, and the evaluation of a large number of gRNA targets. Compared to other cell-based off-target identification methods such as DISCOVER-seq^25–27^, it has the advantage that it captures the range of on- and off-target editing activity that has occurred over time, rather than surveying unrepaired DSBs. It more directly detectes Cas9 nuclease off-target sites, compared to biochemical off-target detection methods such as CIRCLE-seq, CHANGE-seq, and Digenome-seq^55^.

Our novel and streamlined GUIDE-seq-2 approach enabled us to efficiently survey 97% of common genetic variants found with the 1,000 Genomes Project, resulting in 21 high-confidence variants and 13 additional variants with significant effects when measured by GUIDE-seq-2 or targeted sequencing. However, developing a general understanding of how genetic variants influence off-target activity requires high-diversity datasets with even representation of possible mismatches. This gap motivated us to explore alternative solutions that could not be found by genomic off-target profiling.

Massively parallel biochemical profiling with CHANCE-seq has advantages over existing approaches. In contrast to biochemical methods like CHANGE-seq and Digenome-seq, which characterize off-target activity in genomic DNA, CHANCE-seq interrogates large, highly diverse, partially randomized target libraries. These libraries are intrinsically better suited for understanding how different mismatches may affect CRISPR-Cas9 activity and for training AI and machine learning models to predict the impact of genetic variation, because they quantify the impact of every mismatch in every possible position, embedded within millions of diverse sequence contexts.

The largest publicly available, systematically mismatched off-target dataset for model training was generated by Doench et al. While this method evaluates all single-nucleotide mismatches per target, CHANCE-seq expands upon this coverage by roughly 6,000-fold per target and adds many examples of targets with combinatorial mismatches. The CRISPRoffT database also provides a valuable resource with approximately 226,000 off-targets, although its genomic DNA sourcing results in limited per-target coverage.

Most published off-target datasets are enriched for detected cleavage events and lack comprehensive negative examples, i.e.,. those genomic sites that are not recognized or cleaved by the RNP complex. These underrepresented negative data are critically important for robust training of machine learning models and are also captured by CHANCE-seq. Combined with the scale of our dataset, this provides an ideal foundation for training deep learning models. Larger datasets like ours contain the long tail of rare but important exceptional cases, improving models’ ability to recognize subtle sequence patterns and contextual factors, and enhancing performance on new data. This comprehensive data foundation supports more advanced model architectures and may lead to stronger predictive accuracy and reliability across diverse genetic backgrounds and experimental conditions.

Our machine learning approach to predict the biochemical activity of genome editors, CHANCE-net, trained on activity data from millions of homologous target sequences, has notable differences with current state-of-the-art methods. For example, the widely used CFD^56^ score uses a position weight probability matrix to calculate the likelihood of off-target editing activity, where mutations at certain positions are assigned a probability of 1 and thus cannot exert an effect. In contrast, our deep learning approach considers contextual mismatches within an off-target site when making predictions. These approaches enable the estimation of variant effects even when cells from individuals with those specific variants are not readily available. They are also sensitive in unusual cases, such as predicting variants at the far 5’ end of Cas9 targets that still have strong effects.

There are several modifications to our current model that could further enhance its performance in future iterations. Training on a broader, more diverse set of target sites would likely improve generalizability, although we are highly encouraged that training on even a small number of sites with millions of mismatched targets yields a model with state-of-the-art performance. Fine-tuning the model using cellular editing outcomes may better align predictions with biological activity. Finally, incorporating insertions and deletions into off-target libraries will provide a more comprehensive representation of the variant landscape, enabling the model to capture a wider range of genomic alterations that may influence Cas9 activity.

Collectively, our studies represent a solution to defining how individual genetic variation affects genome editing activity for research and therapeutics, that does not require genome-wide activity profiling of cells from a large number of individuals. First, commonly occurring cellular off-target sites could be discovered using methods like GUIDE-seq (or CHANGE-seq) and validated by targeted sequencing, narrowing down choices for highly active and specific targets. Second, complementary CHANCE-seq biochemical profiling of specific lead targets of interest could be performed rapidly to train accurate machine learning predictors to assess population-level risk. These models could be used for hybrid experiment-guided machine-learning prediction to identify highly active and specific gRNA targets that are also least likely to be affected by genetic variation within a subject population. We note that the safety risk associated with off-target mutations is likely to be highly context-dependent, and thus there is no set threshold below which off-target activity is safe and above which it can be tolerated^58^. Nonetheless, variant-aware analysis can help inform a context-dependent risk analysis.

Further research is needed to understand the impact of genetic variation across a larger number of targets and in the context of diverse editor orthologues and types, such as base^60,61^ and prime editors^62^. In the future, with a more generalizable CHANCE-seq model, it may become possible to use the activity learned from biochemical approaches to estimate cellular editing frequencies and to train broadly applicable variant-aware predictors to pick the best highly active and specific genome editing targets for research and therapeutics. Such an approach would shift the reliance on experimental methods from the initial stages of lead target identification to later-stage validation, thereby streamlining the target selection pipeline (**Fig. 6m**).

Overall, we envision a path by which these types of cellular, biochemical, and artificial intelligence approaches could be used to rapidly chart a path for considering genetic variation when designing genome editing strategies. For genome editing therapeutics designed to treat large numbers of patients (such as for sickle cell disease and others), it will be possible to predict which common genetic variants are most likely to result in unwanted cellular off-target activity in association with a particular gRNA target. For genome editing therapies designed to correct a rare mutations, individual genetic variation could be used to estimate patient-specific off-target risk. Finally, for general life sciences research it will be generally useful to predict specific targets that are least likely to exhibit confounding, non-specific activity.

## Data availability

High-throughput sequencing data associated with this manuscript (GUIDE-seq-2, CHANCE-seq, targeted sequencing) are available through NCBI GEO SuperSeries GSE286300.

## Code availability

GUIDE-seq-2 analysis code available through https://github.com/tsailabSJ/guideseq. CHANCE-seq analysis code available through https://github.com/tsailabSJ/chanceseq repository.

## Acknowledgements

We thank the St. Jude Protein Production Core Facility for recombinant Tn5 expression and Sandra Capellera Garcia for scientific editing assistance. This research was supported by the National Institute of Allergy and Infectious Diseases (NIAID) of the National Institutes of Health (NIH) under award numbers U01AI176470 and U01AI176471, St. Jude Children’s Research Hospital and ALSAC, and St Jude Children’s Research Hospital Collaborative Research Consortium on Novel Gene Therapies for Sickle Cell Disease. J.M. received additional support from a Guggenheim Fellowship from the John Simon Guggenheim Memorial Foundation and the Ray and Stephanie Lane Professorship at Carnegie Mellon University. The funders had no role in study design, data collection and analysis, decision to publish or preparation of the manuscript.

## Author contributions

C.R.L., Y.L., J.C, and A.R.F. designed the research, performed experiments, analyzed data, and wrote the manuscript. W.Y. designed and trained the deep learning model. V.K., E.U., G.L., R.W., and A.M. performed human cell experiments. S.R.R. performed variant association analysis. J.M. supervised the deep learning analysis. Y.C. and S.Q.T. designed and supervised the research and wrote the manuscript with input from all authors.

---

**Extended Data Fig. 1.**
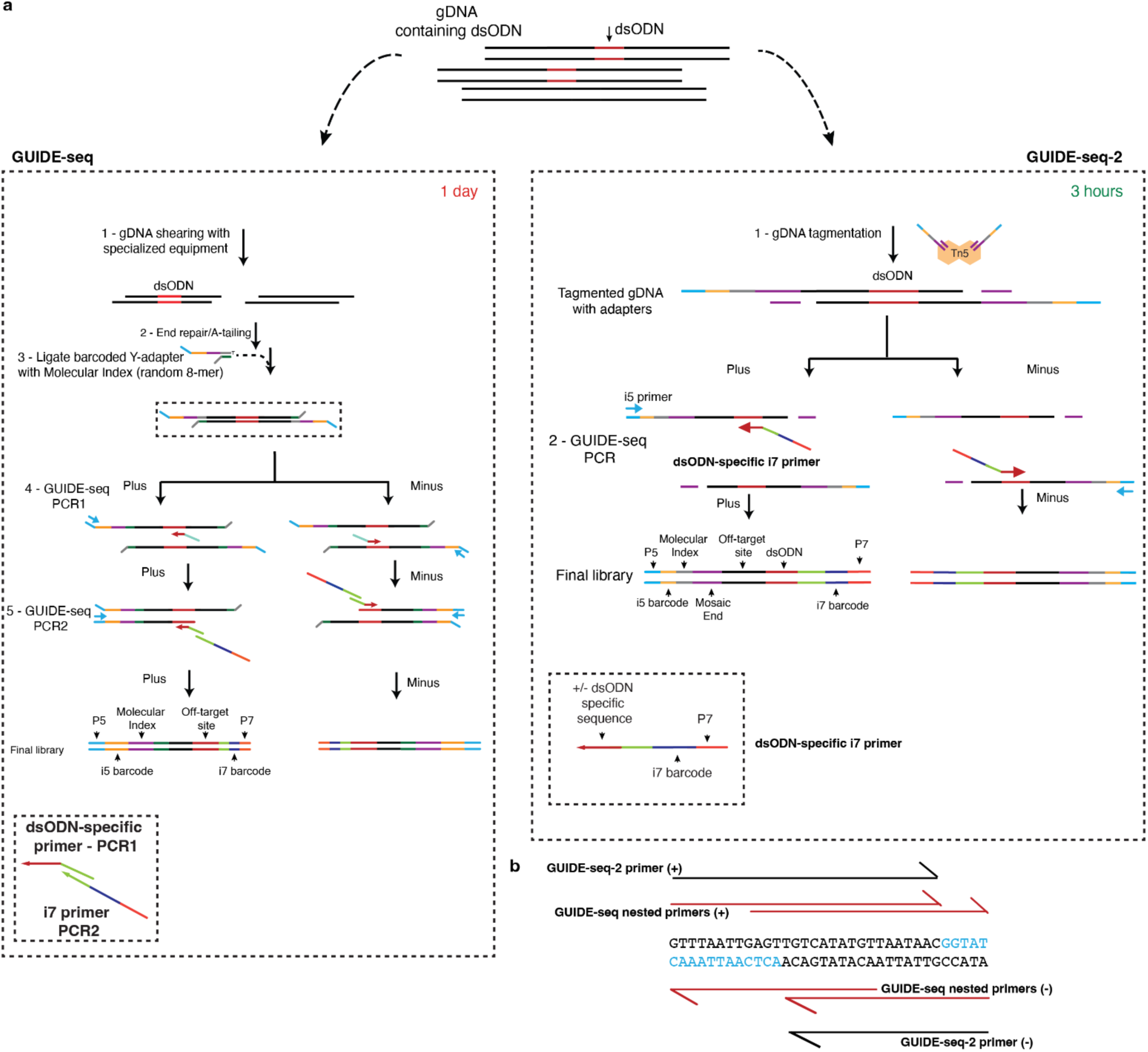
Schematic overview of GUIDE-seq and GUIDE-seq-2. **a,** Both GUIDE-seq and GUIDE-seq-2 begin with cells transfected with a nuclease genome editor and a GUIDE-seq double-stranded oligodeoxynucleotide (**dsODN**) tag. GUIDE-seq requires fragmentation of genomic DNA (typically by physical shearing with sonication) followed by end-repair/A-tailing, ligation, and nested PCR. GUIDE-seq-2 eliminates the requirement for specialized equipment for physical DNA shearing along with 6 additional molecular biology or purification steps by leveraging Tn5 transposase for library preparation and eliminating the requirement for nested PCR. First, genomic DNA from GUIDE-seq dsODN tag integrated cells is tagmented with unique-molecular index containing transposomes. Tagmented gDNA is then subjected to a single round of PCR amplification, using a tag-specific primer combined with the i7 barcode primer, obviating the need for a second round of PCR. **b,** GUIDE-seq and GUIDE-seq-2 forward and reverse tag-specific primers are shown. Use of a single PCR step for library preparation enabled a primer design capable of distinguishing mispriming artifacts without changing the length of GUIDE-seq tag and affecting the tag integration efficiencies. Bases expected to be observed with the new primer design in the GUIDE-seq-2 are highlighted in blue. The simplified GUIDE-seq-2 workflow substantially streamlines the process and enables high-throughput experiments, while also decreasing the requirement of input genomic DNA for library preparation by approximately 4-fold.

**Extended Data Fig. 2.**
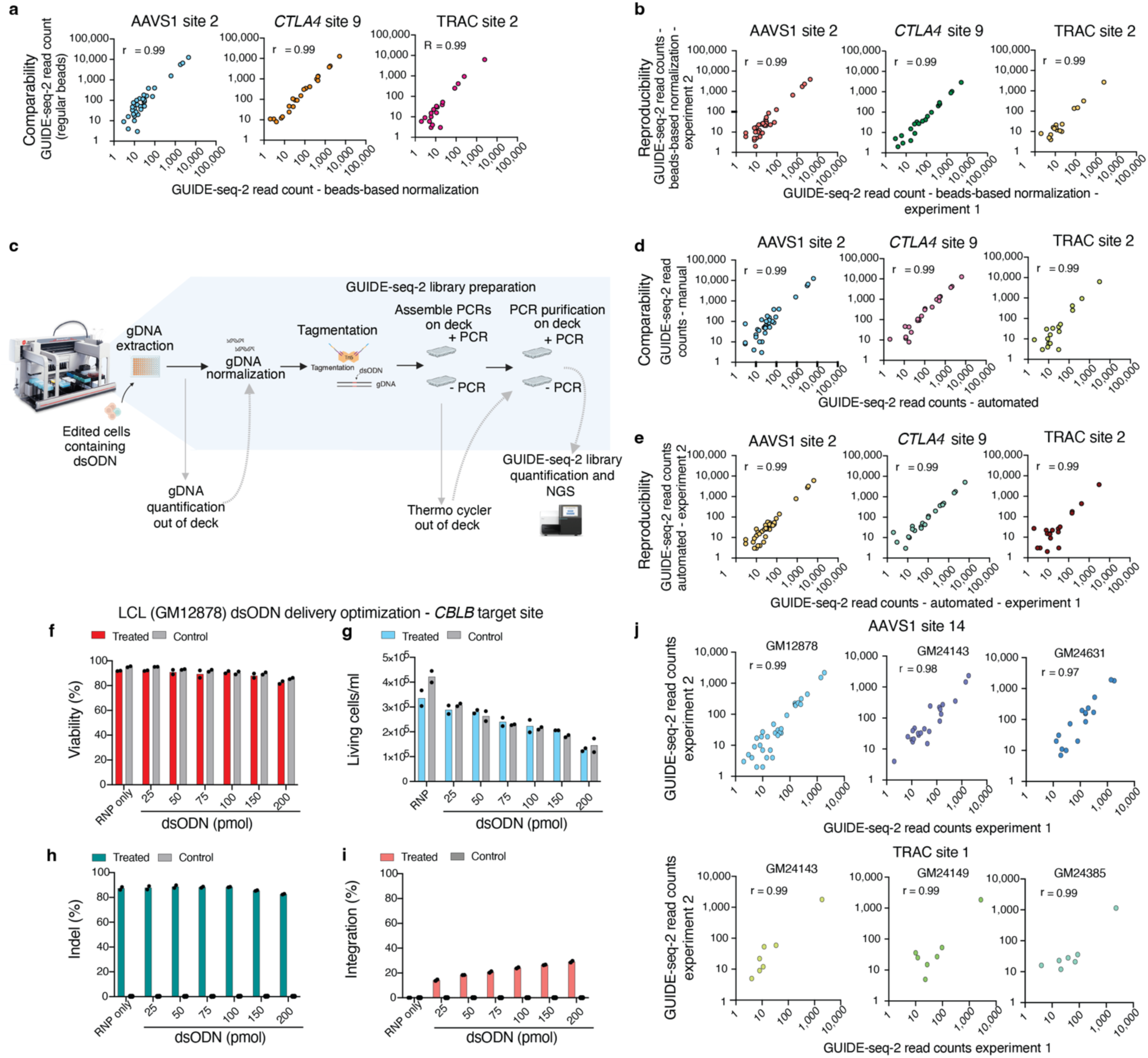
GUIDE-seq-2 automation of library preparation and optimization for lymphoblastoid cell lines. **a,** Scatterplots of GUIDE-seq-2 read counts (log scale) between beads-based library normalization and regular SPRI beads purification. **b,** Scatterplots of GUIDE-seq-2 read counts (log scale) between two independently prepared beads-based library normalization libraries. **c,** Schematic representation of automated GUIDE-seq-2 library preparation workflow. **d,** Scatterplots of GUIDE-seq-2 read counts (log scale) between GUIDE-seq-2 libraries prepared using the automated workflow and the standard manual workflow. **e,** Scatterplots of GUIDE-seq-2 read counts (log scale) between two independently prepared GUIDE-seq-2 libraries using the automated workflow. **f,** Viability and **g,** live cell count, of lymphoblastoid cell line (LCL) population assessed 3 days post nucleofection with different doses of dsODN (n=2). **h,** Indels rates at the intended target site 3 days post nucleofection with different doses of dsODN. **i,** dsODN integration rates 3 days post nucleofection with different doses of dsODN. **j,** Scatterplots of GUIDE-seq-2 read counts (log scale) between two independently prepared GUIDE-seq-2 libraries for two sgRNA target sites across three LCL donors, showing GUIDE-seq-2 technical reproducibility, **r** is Pearson’s correlation coefficient.

**Extended Data Fig. 3.**
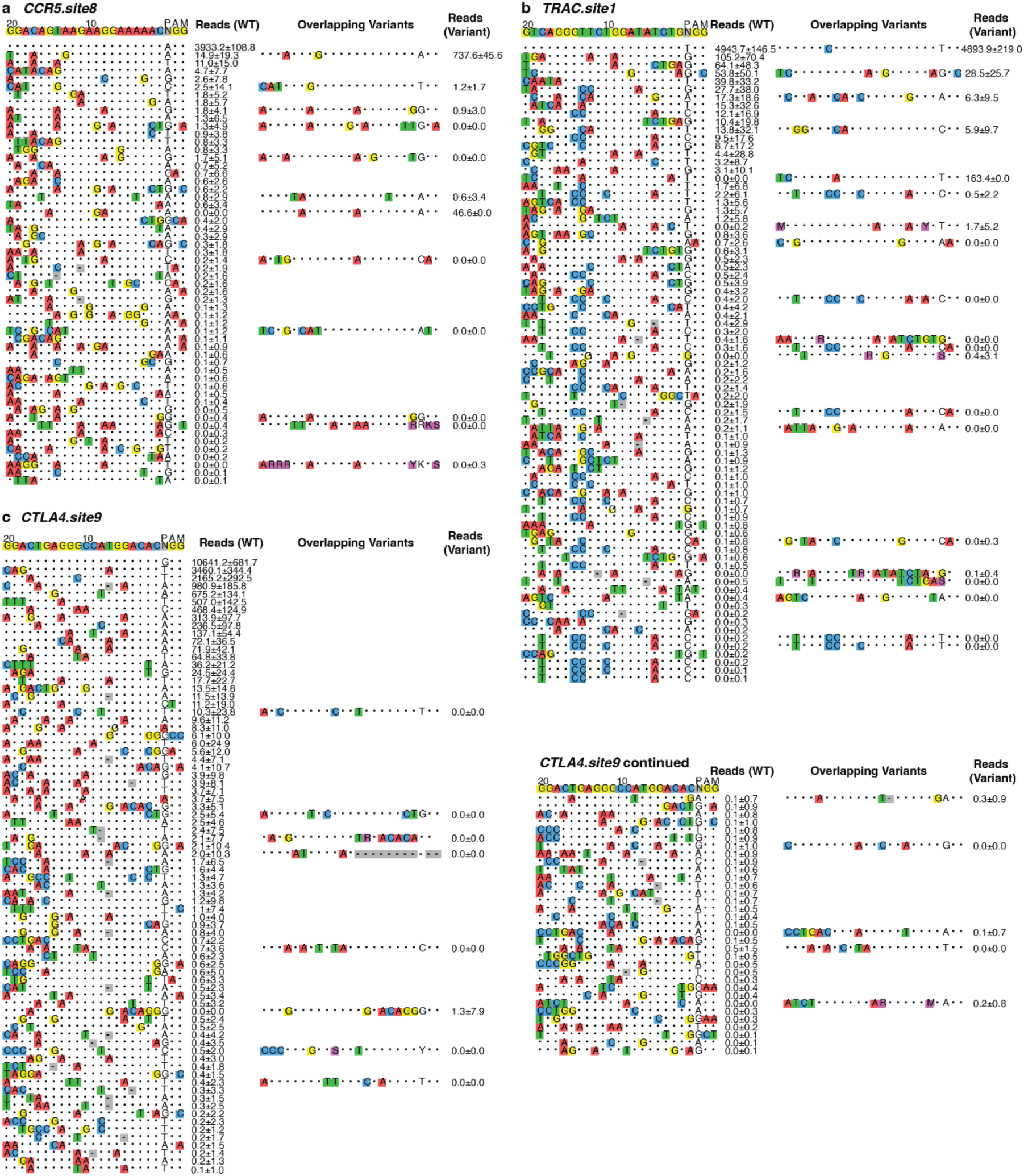
GUIDE-seq-2 visualization plots for 3 targets. The intended on-target site is shown at the top of the alignment visualization with colored nucleotides. Each row shows a genomic off-target site with average of GUIDE-seq-2 read counts and standard deviation (n=95) shown to the right. Mismatches to the intended target site are shown as colored nucleotides, matches are indicated with a dot. The first nucleotide of the NGG PAM sequence is shown without color. The same genomic off-target site but with an overlapping variant sequence is shown further to the right. The average and standard deviation for the variant group (i.e., donors of heterozygous or homozygous alternative genotype) is shown next to it.

**Extended data figure 4.**
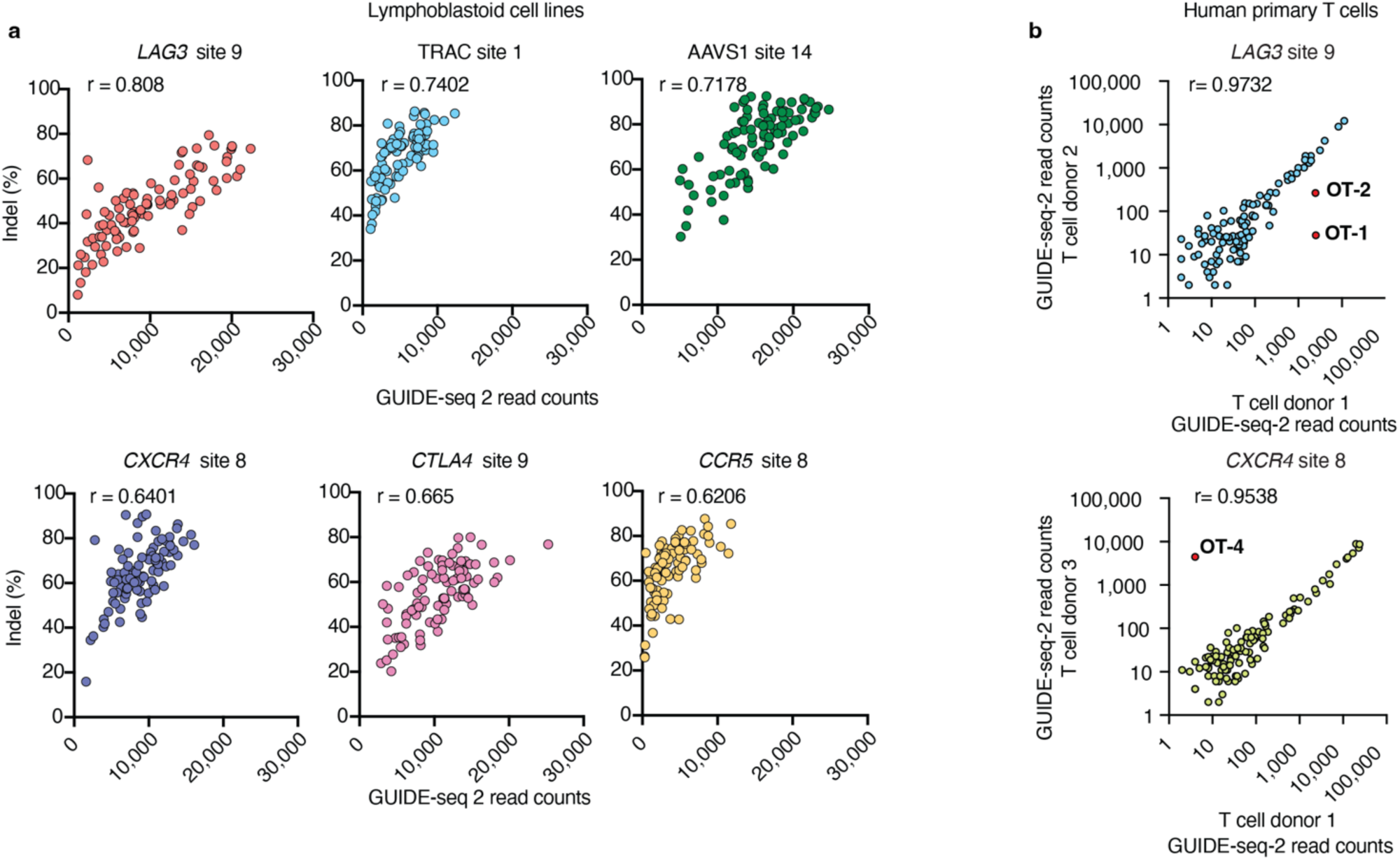
Correlations of GUIDE-seq-2 read counts with indel mutations in LCLs and between donors in human primary T cells. **a,** Correlation of on-target GUIDE-seq-2 read counts and indel (%) for the six sgRNAs target sites evaluated across the 95 lymphoblastoid cell line (LCL) donors, r is Pearson’s correlation coefficient. **b,** Correlation of GUIDE-seq-2 read counts between different human primary T cell donors, dots in red are the examples of off-targets containing SNPs that are visualized in detail in Fig. 4g.

**Extended data figure 5.**
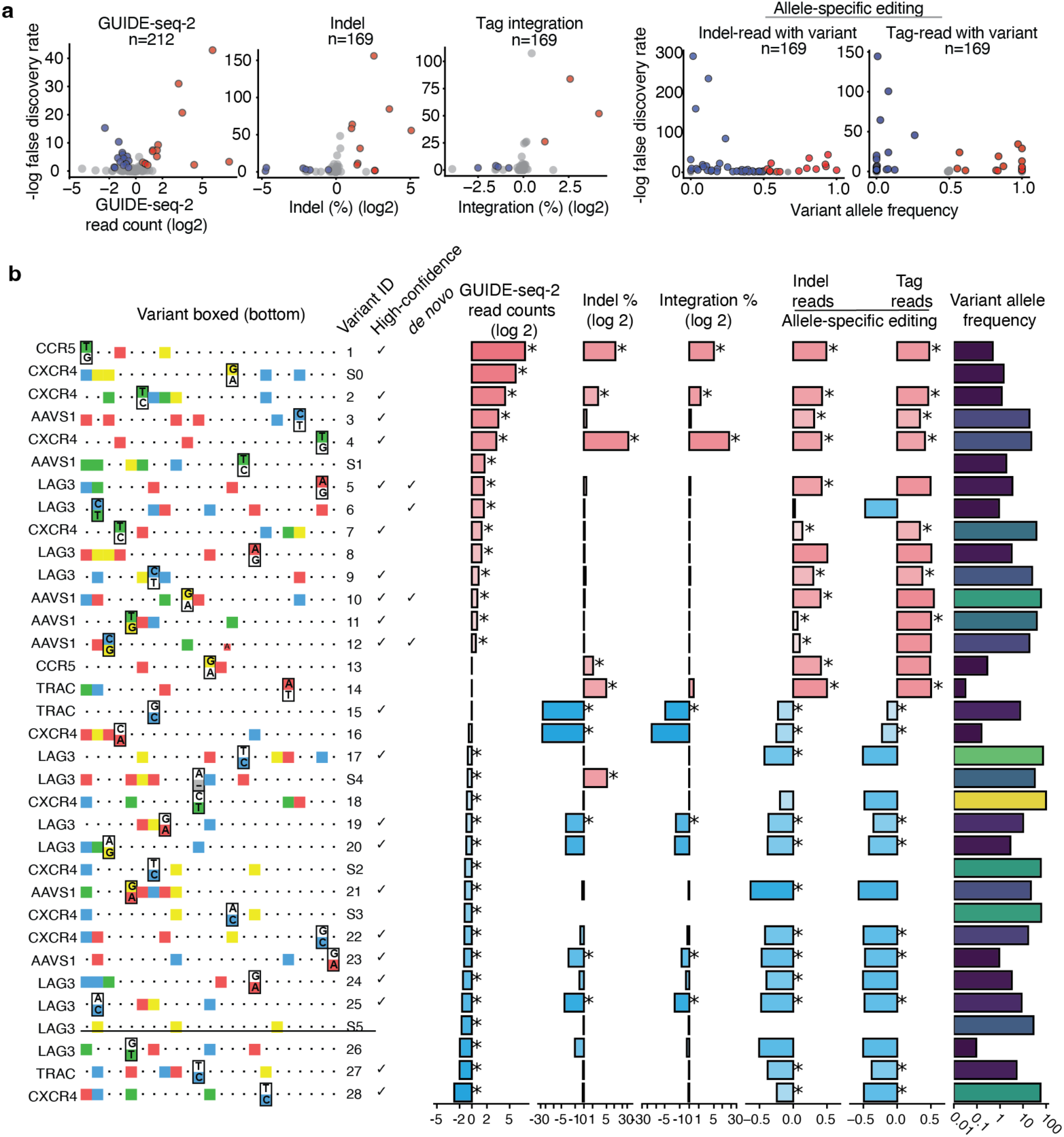
Detailed view of overlapping genetic variants impacting off-target activity identified by GUIDE-seq-2. **a,** Volcano plots showing variant effects on GUIDE-seq-2 read counts, indels, integration, allelic imbalance (x-axis). Y-axis is statistical significance (negative log false discovery rate, FDR). Red dots represent significantly increased variant activity (FDR<=0.1) and blue dots represent significantly decreased variant activity (FDR<=0.1). Absolute difference >=0.5 threshold is used in addition to FDR for GUIDE-seq-2 read counts, indels, and integration. **b,** A detailed view of variants, defined by GUIDE-seq-2 and multiplexed targeted sequencing data, sorted based on GUIDE-seq-2 read counts. Similar to Figure 4f but with variants added only identified by GUIDE-seq-2 and not multiplexed targeted sequencing. Also includes variants with deletions indicated by black lines in the sequence alignment.

**Extended data figure 6.**
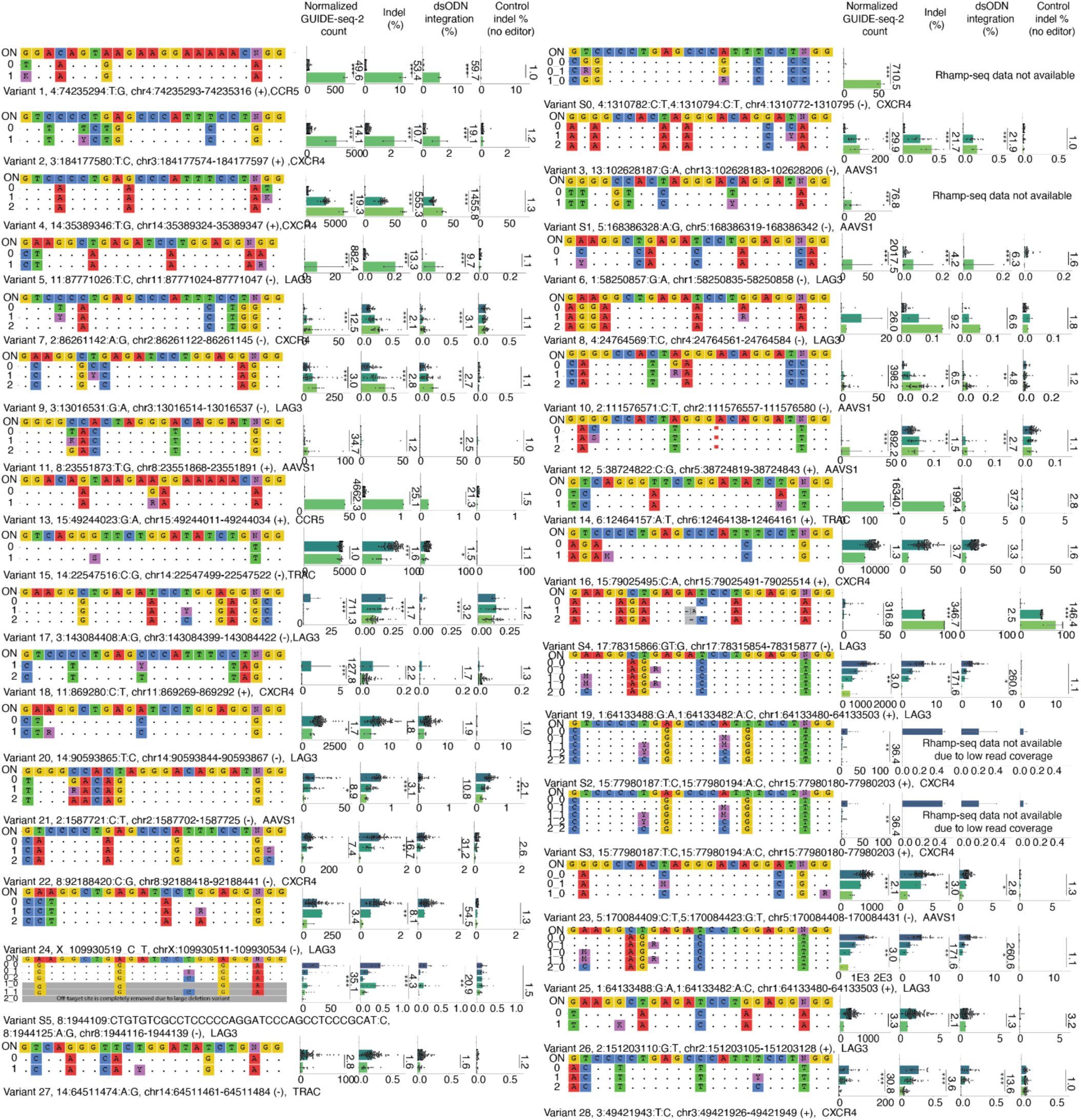
Visualizations of off-targets with variant effects defined by GUIDE-seq-2 and/or multiplexed targeted sequencing. Each off-target visualization panel contains: a sequence-alignment plot, bar plots of GUIDE-seq-2 read counts, indel percentage, dsODN integration percentage, and indel percentage from a no editor control (from left to right). From top to bottom in the sequence-alignment plot shows the on-target sequence (i.e., ON) and the off-targets sequences corresponding to homozygous reference genotype (0), and may contain heterozygous genotype (1), and homozygous alternative genotype (2). Fold change comparing homozygous reference genotype and homozygous alternative genotype is shown. Student’s t-test significance is shown. *p-value<=0.01. **p-value<=0.001. ***p-value<=0.0001. Variant ID S2 and S3 occur in the same off-target, therefore, multiple genotypes exist. IUPAC DNA codes are: K: T or G. M: A or C. S: C or G. Y: C or T. R: A or G. N: A, C, G, or T. Off-target annotation is shown on the bottom, including variant ID (corresponding to figure 4f), variant location (hg38), off-target coordinate, DNA strand (+ or -), and on-target name.

**Extended Data Fig. 7.**
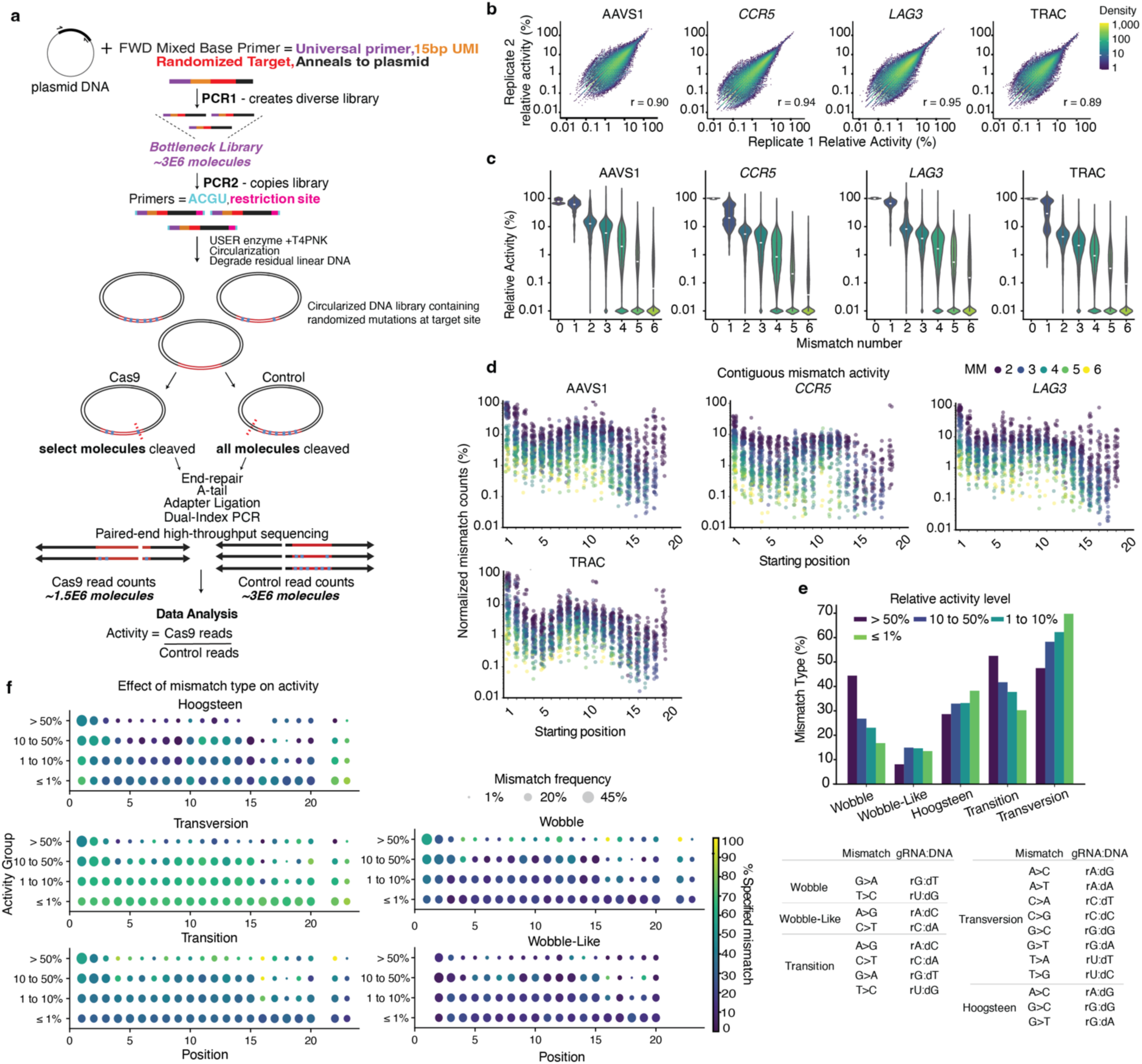
CHANCE-seq detailed schematic and plots. **a,** Detailed schematic of CHANCE-seq assay. Composition of primers are color coded. In the circles, red represents the target site, black is the plasmid backbone, blue squares represent randomized mismatches in protospacer and PAM. PCR2 products are circularized, and the library is then split between Cas9 and restriction enzyme control ensuring the same composition in both samples. Restriction enzyme cleaves all randomized targets, select molecules are cleaved with Cas9. Only cleaved circles are sequenced. **b,** Hexbin plot of relative activity between replicate 1 and replicate 2 for Cas9 CHANCE-seq library using gRNA. Each dot is an off-target. 4 targets are shown colored by density. **c,** Violin plots showing % relative activity at each mismatch level 0-6 for 4 targets, target using CHANCE-seq. **d,** Strip plot illustrating contiguous mismatch start positions (2-6 mismatches) with % relative activity on a log scale. Values plotted for 4 targets. **e,** Bar plot showing effect of mutation type on activity. All CHANCE-seq off-targets were divided into 5 groups based on relative activity. For each activity group, the percentage of off-targets falling into 1 of 5 mismatch categories is shown **f,** Heatmap of mutation types by activity level and protospacer position. Circle size represents frequency of any mismatch at each position within each activity group and color indicates the percentage of specific mismatch type at each position.

**Extended Data Fig. 8.**
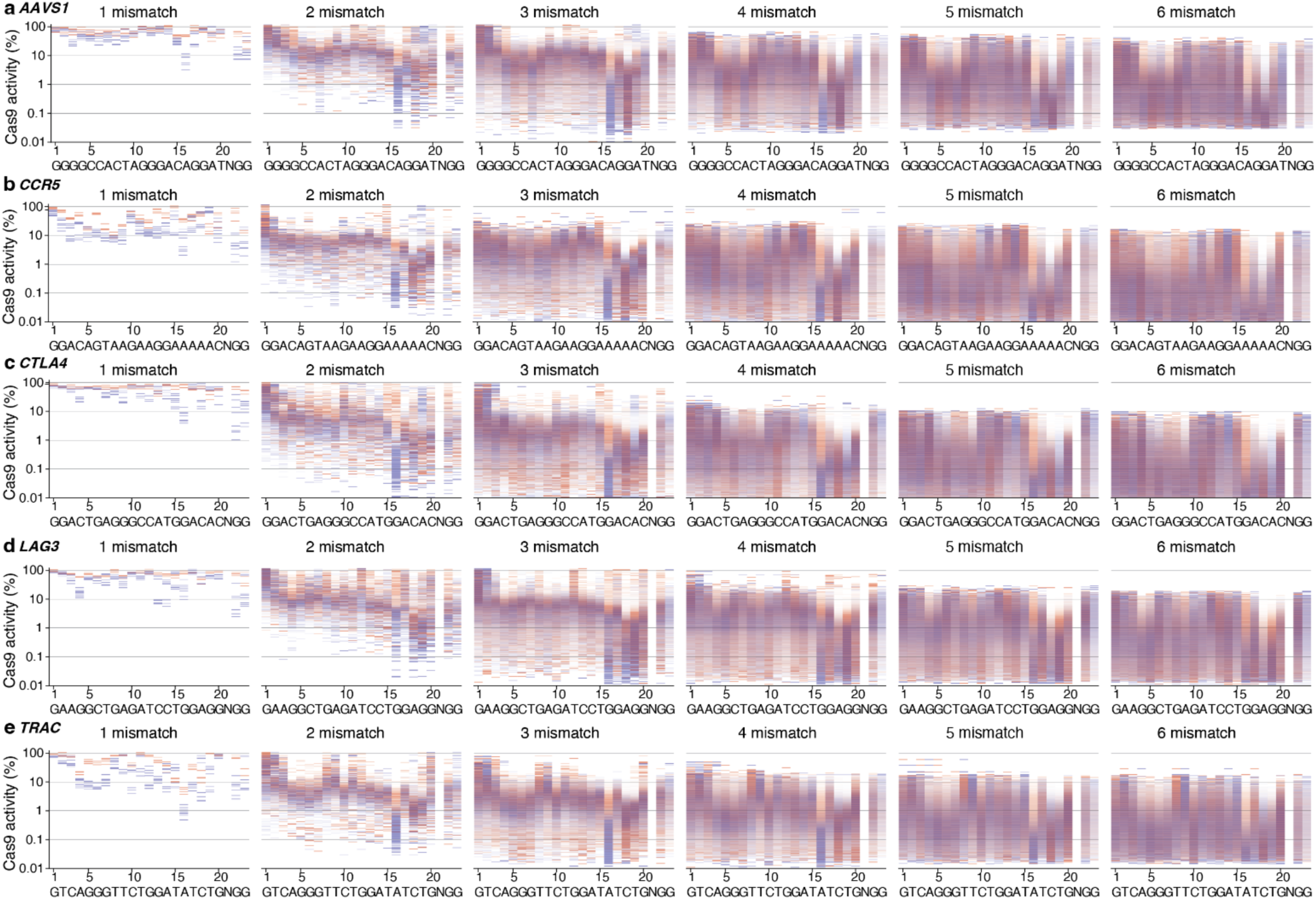
CHANCE-seq global visualizations for 5 targets. Global analysis showing log10 relative activity for **a,** AAVS1 **b,** *CCR5* **c,** *CTLA4*, d,*LAG3* and **e,** TRAC using CHANCE-seq, stratified by mismatch. The average of 2 technical Cas9 replicates was taken to determine relative activity percentage. Plots are divided into bins based on activity. Transitions are colored in red and transversions are in blue, with transparency at each position proportional to position mismatch frequency within each bin. On-target sequence is shown at the bottom of the plot.

**Extended Data Fig. 9.**
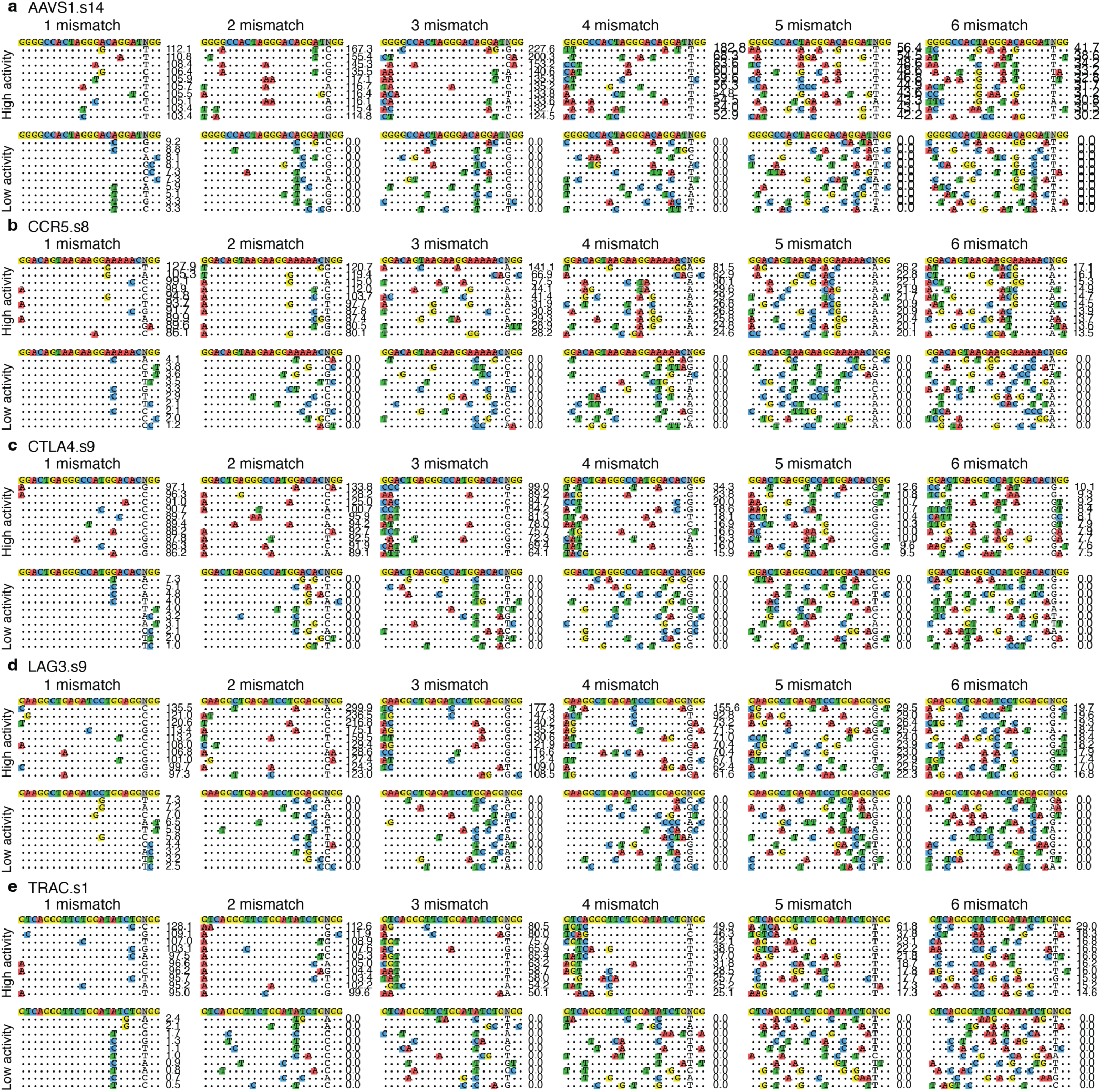
Visualization of sequences that had the top 10 highest and lowest relative activity for. **a,** AAVS1 **b,** *CCR5* **c,** *CTLA4,* **d,** *LAG3* and **e,** TRAC using CHANCE-seq. Sequences are stratified by mismatch number and the average of 2 technical Cas9 replicates was taken to determine relative activity. The top sequence is the on-target, each subsequent sequence shows where mismatches occur. The number to the right shows the percent relative activity for each sequence.

**Extended data figure 10.**
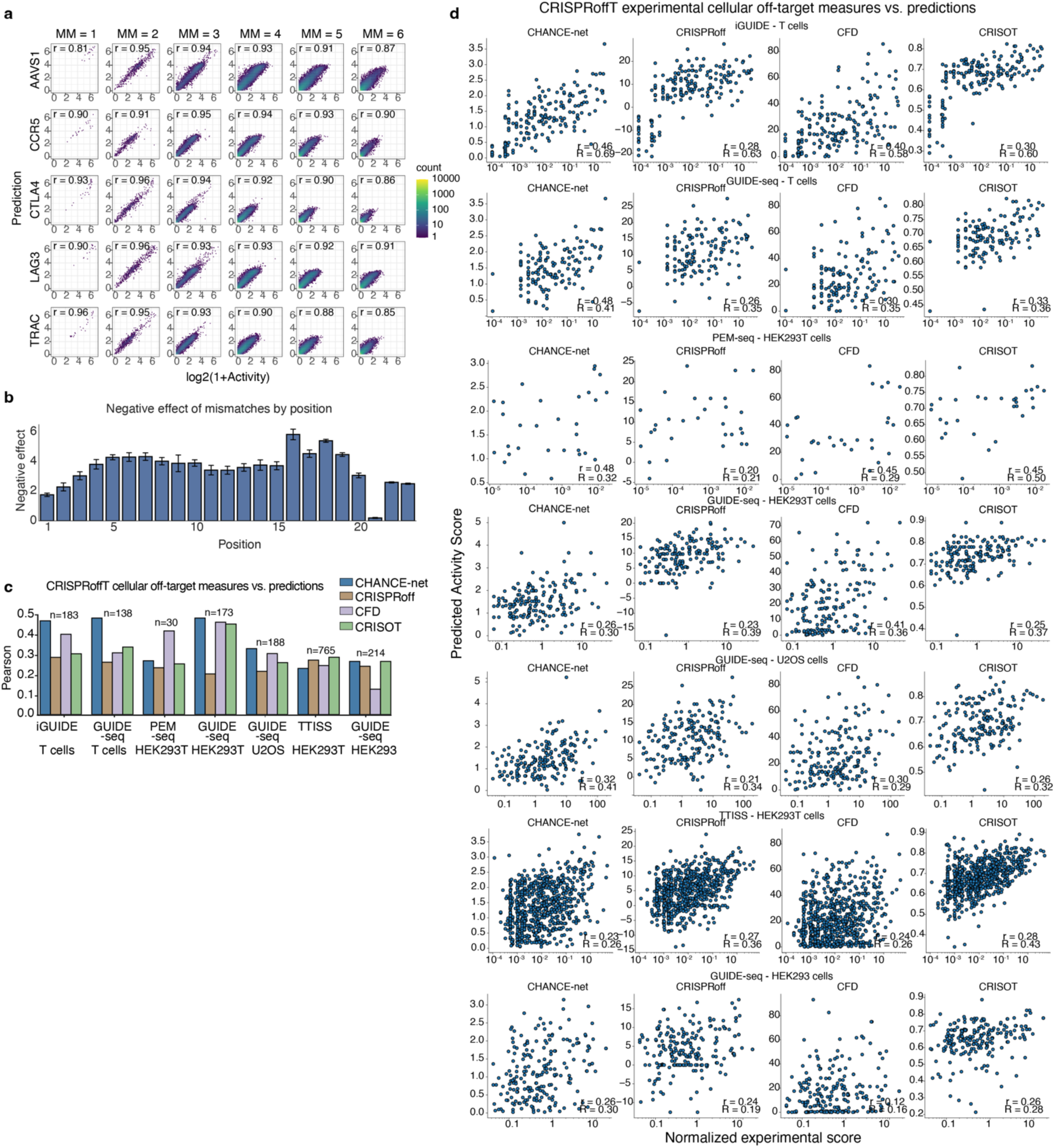
CHANCE-net machine learning performance. **a,** Scatterplot of machine learning performance stratified by mismatch and target site to predict activity within CHANCE-seq datasets. Experimental values are shown on the x-axis and predicted values from 10% hold-out data are on the y-axis, *r* = Pearson’s. **b,** Bar plot of position-wise feature importance scores, represented as the average negative effect of mismatches at each position across all test sequences. y-axis = magnitude of the negative effect, x-axis = sequence position. Error bars = variability across 5 gRNA targets. **c,** Bar plot benchmarking performance of CHANCE-net and 3 other predictive models on CRISPRoffT cellular off-target dataset. Pearson correlation is between predicted off-target activity and experimentally measured off-target activity across seven cell type/assay groups. n = number off-target sequences in each group. **d,** Scatter plots comparing predicted off-target activity (y-axis) with normalized experimental activity (x-axis) across seven cell type/assay groups in the CRISPRoffT dataset. Rows correspond to cell type/assay groups, and columns correspond to prediction methods. Each point represents one off-target sequence. r=Pearson correlation R=Spearman.

**Supplemental Figure 1.**
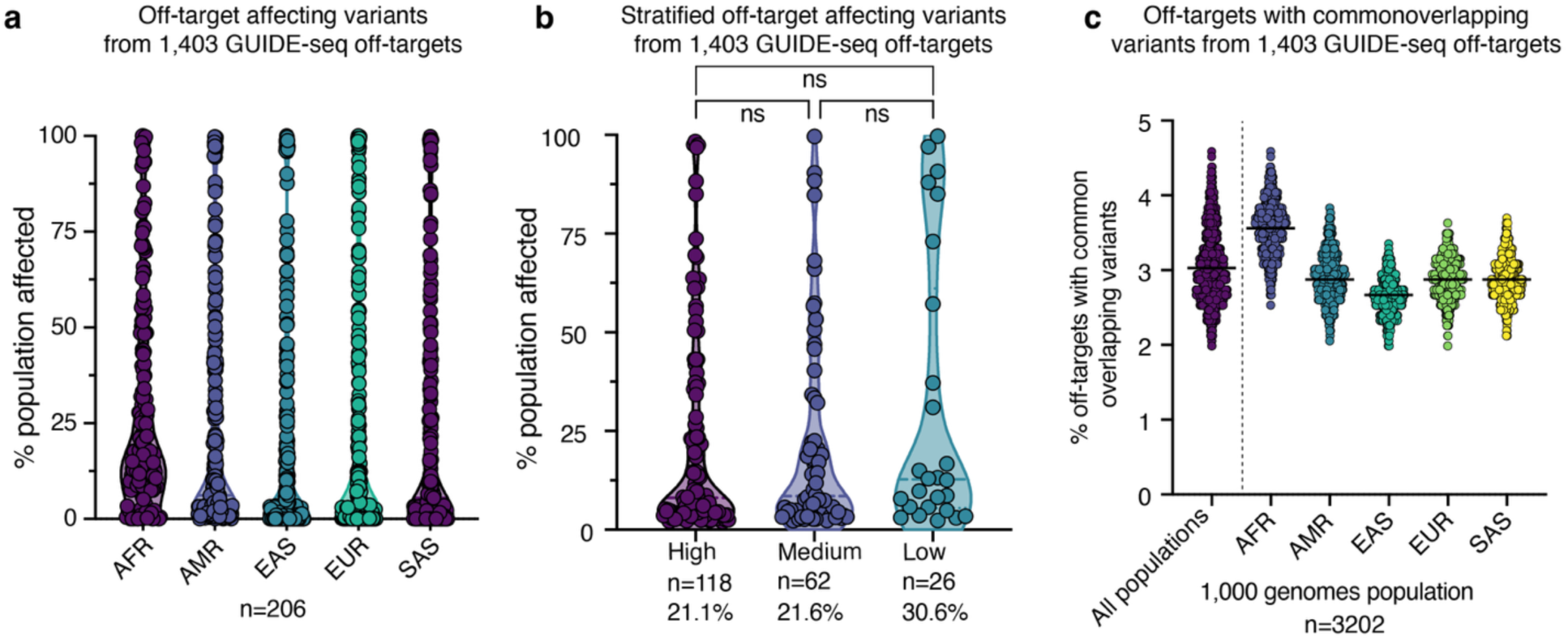
GUIDE-seq identified off-targets applied to 1,000 Genomes Project for variant discovery. **a,** Violin plot showing percentage of population affected by a given off-target affecting variant. Each dot is a variant. **b**, Violin plot showing percentage of population affected stratified by on-target specificity. The three groups are based on the number of off-targets per on-target. Low group contains the most specific on-targets, defined as off-targets per on-target less than 20. The total number of off-target affecting variants is 26. Median group contains on-target with off-targets numbers range from 20 to 200. The total number of off-target affecting variants is 62. High group contains on-target with off-targets numbers more than 200. The total number of off-target affecting variants is 118. Significance determined by 2-way ANOVA. **c**, Violin plot showing percentage of off-targets overlapping with common variants on a per individual basis. All 1,000 genomes population is show on the left, then by separated by superpopulation on the right. African/African American (**AFR**), Admixed American (**AMR**), East Asian (**EAS**), (**EUR**) South Asian (**SAS**). Each dot is a donor from the 1000 Genome Project.

