## Supplementary information for "Frequent, context-dependent effects of human genetic variation on Cas9 activity revealed by population-scale GUIDE-seq-2 and deep combinatorial CHANCE-seq profiling"

#### Supplementary note 1: GUIDE-seq-2 development

GUIDE-seq-2 was developed to overcome limitations of the original GUIDE-seq protocol[1,2], which included a library preparation protocol that relied on a specialized instrument for genomic DNA sonication and shearing, a multi-step library preparation protocol that required end-repair, adapter ligation, A-tailing, and two rounds of tag-specific PCR, library quantification and sequencing with custom primers.

Using a 'less is more' design principle[3] we searched for subtractive changes to the protocol that could simultaneous improve its performance and efficiency. We applied this delete principle to remove the following steps: 1) physical genomic DNA sonication and shearing with a specialized and expensive instrument, 2) end-repair, adapter-ligation, and A-tailing, 3) one round of PCR, and 4) library quantification by qPCR.

Towards that end, we replaced physical genomic DNA shearing with Tn5 tagmentation which enabled simultaneous DNA fragmentation and shearing. We combined primers that had been previously delivered in two parts into one, improving GUIDE-seq-2 chemistry with a PCR design that enabled mispriming detection via observation of expected bases after the annealed PCR primer. We replaced library quantification by qPCR with a streamlined bead-based normalization approach. The new library design also eliminates the earlier GUIDE-seq requirement for addition of custom sequencing primers, which may broaden accessibility of GUIDE-seq to laboratories which do not have access to customizing short-read, high-throughput sequencer conditions. Finally, we carefully tested the final combination of improvements for comparability to original GUIDE-seq in the human primary T cell experiments described in this study.

Tn5 can be purified in the lab as described previously[4] (and summarized below) or purchased commercially. We tested Tagify-UMI obtained from seqWell and found that GUIDE-seq-2 results were strongly correlated as shown in Supplementary Fig. 1.

#### Supplementary protocol 1: GUIDE-seq-2

**Reagents**

- IDTE pH 8.0 (1X TE Solution) (Integrated DNA Technologies, 11050204)
- IDT custom oligonucleotides
- Synthetic chemically modified sgRNA (Synthego)
- Tn5 transposase
- Proteinase K (NEB,P8107S)
- Ultrapure Nuclease Free water (Invitrogen 10977023)
- Magnum FLX magnetic rack (Alpaqua, A00400)
- dNTPs (NEB, cat.no. N0447L)
- Qubit dsDNA HS Assay Kit (Thermo Fisher Scientific, Q32854)
- Lib Quant Kit (Illumina/Uni) (Kapa Biosystems, KK4824)
- Illumina
- PhiX Control V3 KIT (Illumina, FC-110-3001)
- Ultra pure EDTA 0.5 M (Invitrogen, 15575038)
- Ethanol (Sigma, cat.no. E7023)
- Sera-Mag Magnetic Beads; Carboxyl, Speedbeads; hydrophobic; 5 solids (Fisher/GE,. 9981123)
- Guanidine thiocyanate (Sigma, G9277)
- Sodium Chloride 5 M Sterile (Invitrogen, AM9760G)
- TRIS Buffer 1.0 M solution, pH 8.0 (Fisher, 50146868)
- Polyethylene Glycol 8000 (Fisher, 507516674)
- 1M Magnesium chloride (Invitrogen, AM9530G)
- Tween-20 (Sigma, P7949-500ML)
- Sodium Hydroxide solution (Sigma, 72068-100ML)

**Reagent setup**

**SPRI-guanidine binding buffer.** SPRI-guanidine binding buffer is composed by 4 M guanidine thiocyanate, 40 mM TRIS, 17.6 mM EDTA, pH 8.0. To prepare 100 ml of SPRI-guanidine binding buffer, weight 47.26 g of guanidine thiocyanate, add water and keep it on a magnetic stirrer. Once guanidine thiocyanate is dissolved, add 4 ml of TRIS 1 M pH 8 and 3.52 ml of EDTA 0.5 M pH 8. Bring the pH to 8 with HCl.

**Sera-Mag Magnetic Beads preparation.** Add 1 ml of Sera-Mag Magnetic Beads (Fisher/GE) to a 1.5 ml Eppendorf tube. Place in a magnetic rack. Remove the liquid. Remove the tube from the rack. Add 1 ml of TE and homogenize. Place back in the magnetic rack and remove the liquid. Repeat this step for a total of two TE pH 8.0 washes. Then, add 1 ml of TE pH 8.0. Note: this beads preparation step is required for preparing SPRI-RNA beads, SPRI-DNA beads and SPRI-guanidine beads.

**SPRI-guanidine beads preparation.** Add 10 ml of 5 M NaCl to 9 g of PEG 8000 and then add SPRI-guanidine binding buffer (prepared as described above) up to 49 ml. Homogenize during 5 min. Add 1 ml of Sera-Mag Magnetic Beads in TE (prepared as described above) and homogenize. Keep at 4 °C. SPRI-guanidine beads can be stored at 4°C for up to 6 months.

**SPRI-DNA beads preparation.** Add 10 ml of 5 M NaCl, 500 µl of 1 M TRIS and 100 µl of 0.5 M EDTA to 9 g of PEG 8000. Complete the volume to 49 ml with ultra-pure water. Add 1 ml of Sera-Mag Magnetic Beads in TE (prepared as described above) and homogenize. Add 27.5 µl of Tween-20 and homogenize. Keep at 4°C. SPRI- beads can be stored at 4 °C for up to 6 months. SPRI-DNA beads can be replaced by AMPure XP beads.

**Tn5**

pTXB1-Tn5 (addgene # 60240) was expressed in Rosetta(DE3)pLysS (millipore sigma #71403). After cell lysis and clarification, Tn5 was purified with chitin beads on HiPrep Q Sepharose 16/10 column (Cytiva) and cleaved with 20 mM HEPES, pH 7.2; 200 mM NaCl, 1 mM EDTA, 10 % glycerol, 0.2 % Triton X-100, 50 mM DTT. Eluted Tn5 was further purified with size exclusion and dialyzed into 100 mM HEPES pH7.2, 200 mM NaCl, 0.2 mM EDTA, 2 mM TCEP, 0.2% TritonX-100, 20% glycerol. To make stocks, purified Tn5 was diluted to 4.5 mg/mL with 2x Tn5 buffer (below) and further mixed with Glycerol and 2x Tn5 buffer at 1 : 1.1 : 0.33 to make 1.85 mg/mL Tn5 final. Stocks were stored at -20C until transposome assembly.

**2X Tn5 dialysis buffer**

To prepare 500 ml of 2X Tn5 dialysis buffer add as follows:

| Component | Volume (ml) | Final Concentration |
|---|---|---|
| HEPES-KOH (1 M) pH 7.2 | 50 | 100 mM |
| NaCl (5 M) | 20 | 200 mM |
| EDTA (0.5 M) | 0.2 | 0.2 mM |
| DTT (1 M) | 1 | 2 mM |
| Triton X-100 | 1 | 0.2% (vol/vol) |
| Glycerol | 100 | 20% (vol/vol) |
| Nuclease-free water | to 500 ml | |
| Total | 500 | |

2X Tn5 dialysis buffer can be stored at 4 °C for up to 6 months.

**5X TAPS-DMF buffer**: 50 mM TAPS-NaOH pH 8.5, 25 mM $MgCl_2$, 50% (vol/vol) DMF. *Prepare fresh.*

| Component | Volume (µl) | Final Concentration |
|---|---|---|
| TAPS-NaOH (0.5 M) | 100 | 50 mM |
| $MgCl_2$ (1 M) | 25 | 25 mM |
| DMF | 500 | 50% (vol/vol) |
| Nuclease-free water | 375 | |
| Total | 1000 | |

**Primers and oligos**

I5 oligos

| Oligo name | Sequence |
|---|---|
| Tn5-A bottom | /5Phos/CTGTCTCTTATACA/3ddC/ |
| i5_N501_UMI_Tn5-A | AATGATACGGCGACCACCGAGATCTACACTAGATCGCNNNNNNNNNNTCGTCGGCAGCGTCAGATGTGTATAAGAGACAG |
| i5_N502_UMI_Tn5-A | AATGATACGGCGACCACCGAGATCTACACCTCTCTATNNNNNNNNNNTCGTCGGCAGCGTCAGATGTGTATAAGAGACAG |
| i5_N503_UMI_Tn5-A | AATGATACGGCGACCACCGAGATCTACACTATCCTCTNNNNNNNNNNTCGTCGGCAGCGTCAGATGTGTATAAGAGACAG |
| i5_N504_UMI_Tn5-A | AATGATACGGCGACCACCGAGATCTACACAGAGTAGANNNNNNNNNNTCGTCGGCAGCGTCAGATGTGTATAAGAGACAG |

| | |
|---|---|
| i5_N505_UMI_ Tn5-A | AATGATACGGCGACCACCGAGATCTACACGTAAGGAGNNNNNNNNNN TCGTCGGCAGCGTCAGATGTGTATAAGAGACAG |
| i5_N506_UMI_ Tn5-A | AATGATACGGCGACCACCGAGATCTACACACTGCATANNNNNNNNNN TCGTCGGCAGCGTCAGATGTGTATAAGAGACAG |
| i5_N507_UMI_ Tn5-A | AATGATACGGCGACCACCGAGATCTACACAAGGAGTANNNNNNNNNN TCGTCGGCAGCGTCAGATGTGTATAAGAGACAG |
| i5_N508_UMI_ Tn5-A | AATGATACGGCGACCACCGAGATCTACACCTAAGCCTNNNNNNNNNN TCGTCGGCAGCGTCAGATGTGTATAAGAGACAG |

Primers for PCR

| **Oligo name** | **Sequence** |
|---|---|
| Common i5 primer | AATGATACGGCGACCACCGAGATCTACAC |

i7 dsODN primers

| | | |
|---|---|---|
| oML89 | i7_N701_dsODN_+ | CAAGCAGAAGACGGCATACGAGATAGCGGAATGTG ACTGGAGTTCAGACGTGTGCTCTTCCGATCTATACC GTTATTAACATATGACA |
| oML90 | i7_N702_dsODN_+ | CAAGCAGAAGACGGCATACGAGATGATCATGCGTG ACTGGAGTTCAGACGTGTGCTCTTCCGATCTATACC GTTATTAACATATGACA |
| oML91 | i7_N703_dsODN_+ | CAAGCAGAAGACGGCATACGAGATAAGACGGAGTG ACTGGAGTTCAGACGTGTGCTCTTCCGATCTATACC GTTATTAACATATGACA |
| oML95 | i7_N704_dsODN_+ | CAAGCAGAAGACGGCATACGAGATCGAGTCCTGTG ACTGGAGTTCAGACGTGTGCTCTTCCGATCTATACC GTTATTAACATATGACA |
| oML96 | i7_N705_dsODN_+ | CAAGCAGAAGACGGCATACGAGATTCCTCAGGGTG ACTGGAGTTCAGACGTGTGCTCTTCCGATCTATACC GTTATTAACATATGACA |
| oML97 | i7_N706_dsODN_+ | CAAGCAGAAGACGGCATACGAGATGTACGGATGTG ACTGGAGTTCAGACGTGTGCTCTTCCGATCTATACC GTTATTAACATATGACA |
| oML98 | i7_N707_dsODN_+ | CAAGCAGAAGACGGCATACGAGATCATCTCTCGTG ACTGGAGTTCAGACGTGTGCTCTTCCGATCTATACC GTTATTAACATATGACA |
| oML99 | i7_N708_dsODN_+ | CAAGCAGAAGACGGCATACGAGATGTCGGAGCGTG ACTGGAGTTCAGACGTGTGCTCTTCCGATCTATACC GTTATTAACATATGACA |
| oML100 | i7_N709_dsODN_+ | CAAGCAGAAGACGGCATACGAGATACGGAGAAGTG ACTGGAGTTCAGACGTGTGCTCTTCCGATCTATACC GTTATTAACATATGACA |
| oML101 | i7_N710_dsODN_+ | CAAGCAGAAGACGGCATACGAGATAGGAGATGGTG ACTGGAGTTCAGACGTGTGCTCTTCCGATCTATACC GTTATTAACATATGACA |
| oML102 | i7_N711_dsODN_+ | CAAGCAGAAGACGGCATACGAGATAGTACTCGGTG ACTGGAGTTCAGACGTGTGCTCTTCCGATCTATACC GTTATTAACATATGACA |
| oML103 | i7_N712_dsODN_+ | CAAGCAGAAGACGGCATACGAGATGGACTCTAGTG ACTGGAGTTCAGACGTGTGCTCTTCCGATCTATACC GTTATTAACATATGACA |

| | | |
|---|---|---|
| oML92 | i7_N701_dsODN_- | CAAGCAGAAGACGGCATACGAGATAGCGGAATGTG ACTGGAGTTCAGACGTGTGCTCTTCCGATCTGTTTA ATTGAGTTGTCATATGTTAATAAC |
| oML93 | i7_N702_dsODN_- | CAAGCAGAAGACGGCATACGAGATGATCATGCGTG ACTGGAGTTCAGACGTGTGCTCTTCCGATCTGTTTA ATTGAGTTGTCATATGTTAATAAC |
| oML94 | i7_N703_dsODN_- | CAAGCAGAAGACGGCATACGAGATAAGACGGAGTG ACTGGAGTTCAGACGTGTGCTCTTCCGATCTGTTTA ATTGAGTTGTCATATGTTAATAAC |
| oML104 | i7_N704_dsODN_- | CAAGCAGAAGACGGCATACGAGATCGAGTCCTGTG ACTGGAGTTCAGACGTGTGCTCTTCCGATCTGTTTA ATTGAGTTGTCATATGTTAATAAC |
| oML105 | i7_N705_dsODN_- | CAAGCAGAAGACGGCATACGAGATTCCTCAGGGTG ACTGGAGTTCAGACGTGTGCTCTTCCGATCTGTTTA ATTGAGTTGTCATATGTTAATAAC |
| oML106 | i7_N706_dsODN_- | CAAGCAGAAGACGGCATACGAGATGTACGGATGTG ACTGGAGTTCAGACGTGTGCTCTTCCGATCTGTTTA ATTGAGTTGTCATATGTTAATAAC |
| oML107 | i7_N707_dsODN_- | CAAGCAGAAGACGGCATACGAGATCATCTCTCGTG ACTGGAGTTCAGACGTGTGCTCTTCCGATCTGTTTA ATTGAGTTGTCATATGTTAATAAC |
| oML108 | i7_N708_dsODN_- | CAAGCAGAAGACGGCATACGAGATGTCGGAGCGTG ACTGGAGTTCAGACGTGTGCTCTTCCGATCTGTTTA ATTGAGTTGTCATATGTTAATAAC |
| oML109 | i7_N709_dsODN_- | CAAGCAGAAGACGGCATACGAGATACGGAGAAGTG ACTGGAGTTCAGACGTGTGCTCTTCCGATCTGTTTA ATTGAGTTGTCATATGTTAATAAC |
| oML110 | i7_N710_dsODN_- | CAAGCAGAAGACGGCATACGAGATAGGAGATGGTG ACTGGAGTTCAGACGTGTGCTCTTCCGATCTGTTTA ATTGAGTTGTCATATGTTAATAAC |
| oML111 | i7_N711_dsODN_- | CAAGCAGAAGACGGCATACGAGATAGTACTCGGTG ACTGGAGTTCAGACGTGTGCTCTTCCGATCTGTTTA ATTGAGTTGTCATATGTTAATAAC |
| oML112 | i7_N712_dsODN_- | CAAGCAGAAGACGGCATACGAGATGGACTCTAGTG ACTGGAGTTCAGACGTGTGCTCTTCCGATCTGTTTA ATTGAGTTGTCATATGTTAATAAC |

**Protocol:**

1) **Tn5 oligos annealing**
Resuspend oligos in IDTE pH 8, to 100 µM.
Prepare i5 oligos:

| Component | | Amount (µl) |
|---|---|---|
| Top oligo (100 µM) | i5_N5##_UMI_Tn5-A | 50 |
| Bottom oligo (100 µM) | Tn5-A bottom | 50 |
| **Total** | | **100** |

- Each top i5 oligo is paired with the same bottom oligo (Tn5-A-bottom)

- On a thermocycler, set up the follow annealing program: 95 °C for 5 min, -1 °C/30 seconds for 70 cycles, hold at 4 °C. After annealing, add 100 µl of IDTE pH 8.0 to bring the concentration of the annealed oligonucleotides to 25 µM. Keep the annealed oligonucleotides at -20 °C. The annealed adapters will be used for transposome assembly.

**2) Transposome assembly**

Prepare Tn5 complex with each annealed i5 oligo:

| Component | Amount (µl) |
|---|---|
| Tn5 (1.85 mg/mL) | 36 |
| Pre-annealed oligo (25 µM) | 15 |
| 2X Tn5 dialysis buffer | 52 |
| **Total** | **103** |

- Incubate at RT for 1 hour.
- Store assembled Tn5 at -20 °C.

**3) Tagmentation**

Prepare a tagmentation reaction for each gDNA sample. 100 ng of gDNA is tagmented with Tn5 complexed with a unique i5 barcode.

| Component | Amount (µl) |
|---|---|
| 5x TAPS-DMF | 8 |
| Assembled Tn5 | 2 |
| Nuclease-free water | 10 |
| gDNA (5ng/µL) | 20 |
| **Total** | **40** |

- Incubate the reaction at 55 °C for 7 minutes.
- Dilute proteinase K (NEB) 1:1 in water (2.5 µl of proteinase K and 2.5 µl of water) and add 5 µl of the dilution to tagmentation reaction. Incubate at 55 °C for 15 minutes.
- Purify with 1.8X (81 µL) of SPRI-Guanidine beads, elute in 11 µL of IDTE.

**4) GUIDE-seq-2 PCR**

Prepare PCR reaction. Each tagmented gDNA is split into + and – group (5 µL of input each). Each + and – sample will contain the same i7 barcode, the only difference is that primers will be specific for + and – group.

For the i7 dsODN primers, make a plate with 5 µM of the primers aliquoted. Example is shown below.

| Component | Amount (µl) |
|---|---|
| Nuclease-free H2O | 15.9 |
| 10x Taq polymerase buffer (Mg free) | 3 |
| 10mM dNTP | 0.6 |
| 50mM $MgCl_2$ | 1.2 |
| Platinum Taq polymerase | 0.3 |
| i7 dsODN primer +/- (5µM) | 2 |
| TMAC (0.5M) | 1.5 |
| Common i5 primer (10 µM) | 0.5 |

St. Jude - Confidential

| | |
|---|---|
| Tagmented DNA | 5 |
| **Total** | **30** |

Run the PCR thermocycler program as follows (70 °C Δ-1.0C/cycle), for 15 cycles decrease the temperature for one degree per cycle starting with 70°C:

| Step | Temperature | Time |
|---|---|---|
| Denaturation | 95°C | 5 min |
| 15 cycles: | 95°C<br>70°C, Δ-1$^{0}$C/cycle<br>72°C | 30 s<br>2 min<br>30 s |
| 20 cycles: | 95°C<br>55°C<br>72°C | 30 s<br>1 min<br>30 s |
| extension | 72°C | 5 min |
| End | 4°C | Hold |

- Purify with 1.8X (54 µl) SPRI beads, elute in 30 µl of TE.

**5) qPCR library quantification**

- Make 1:10 dilution of each + and – library from $10^{-1}$ to $10^{-5}$ with IDTE in total volume of 50 µl.
- $10^{-5}$ diluted samples will be quantified in duplicates.

Prepare qPCR master mix:

| Component | Amount (µl) |
|---|---|
| SYBR fast MM + Primer Premix | 12 |
| Nuclease-free water | 4 |
| **Total** | **16** |

Set up qPCR reaction:

| Component | Amount (µl) |
|---|---|
| qPCR master mix | 16 |
| Sample/Standard/nuclease-free $H_2O$ | 4 |
| **Total** | **20** |

- Pool library to achieve 8 x $10^9$ molecules in 5 µl or to 2 nM for NextSeq2000

**6) Sequencing**

Sequence based on the information listed below specific to each sequencer. Index2 is 18nt:

**MiniSeq**: 146, 8, 18, 146
50 µl of denatured library + 6uL of 20 pM PhiX + 444 µl of buffer

**MiSeq**: 146, 8, 18, 146 (GUIDEseq_v2 sample sheet)
10 µl of denatured library + 100 µl of 12.5 pM PhiX + 940 µl of buffer

**NextSeq 550**: 146, 8, 18, 146
120 µl of denatured library + 15 µl of 20 pM PhiX + 1165 µl of buffer

**NextSeq 2000**: 146, 8, 18, 146
550 pM of denatured library + 20% (vol/vol) of 550 pM PhiX (Load 20 uL)

**Tagmentation protocol using commercially available Tn5**
Pre-assembled Tagify™ i5 UMI (seqWell 301210 or 301230) can be used in replacement of the above steps 1) Tn5 oligos annealing through 3) Tagmentation with the following protocol.

**Reagents**
Pre-assembled Tn5: Tagify™ i5 UMI (seqWell 301210 or 301230) custom ordered for custom volume packaging.
3x coding buffer: seqWell 101000
X solution: seqWell 101001
MAGwise Magnetic Purification Beads: seqWell 101003

**Tagmentation with Tagify™ i5 UMI**
Prepare tagmentation reaction for each gDNA sample. 100 ng of gDNA is tagmented with Tn5 complexed with a unique i5 barcode.

| Component | Amount (µl) |
|---|---|
| 3x Coding buffer | 13.3 |
| Assembled Tn5 | 4 |
| Nuclease-free water | 2.7 |
| gDNA (5ng/µL) | 20 |
| **Total** | **40** |

- Incubate the reaction at 55 °C for 7 minutes.
- Add 20 µl of X solution to the 40 uL tagmentation reaction. Incubate at 68 °C for 10 minutes.

Purify with 1.8X (108 µL) of MAGwise beads and perform the wash/elution step the same as above

### Supplementary note 2: CHANGE-seq-R development

**CHANGE-Seq R optimization**

**End-repair.** After cleavage of the circularized, partially-randomized libraries with RNP complex, library preparation of resulting linear DNA was initially performed without end-repair. Consequently, data analysis showed unexpected enrichment profiles. Since the library is comprised of large numbers of off-targets with a range of mismatches, we hypothesized that staggered cleavage was likely occurring frequently for some targets, resulting in 5' overhangs and therefore, end-repair was necessary to observe the full spectrum of cleavage outcomes during library preparation for Illumina sequencing.

**Assay Development.** Our initial experiments for CHANGE-Seq R utilized primers that did not have a unique molecular index (UMI) incorporated next to the randomized target site. We also used only one PCR to create a diverse library and then produced the circularized library from the products of this PCR. Without an initial bottlenecking step, this resulted in a library with hundreds of millions of unique molecules which made it challenging to obtain adequate sequencing depth in our control samples. Because there was not an added PCR step to make copies of the library, many off-targets that were present in the Cas9 cleaved were not present in the control sample and vice versa. Additionally, the end repair process resulted in base insertions in some Cas9 cleaved off-targets but not in control (SrfI produces blunt ends). Because there were no UMIs in the primers, this made it difficult to confidently locate the same off-target sequence in both the Cas9 and control sample and therefore difficult to calculate enrichment of a particular off-target. To overcome these issues, we added 15 bp UMIs to our primers and an additional PCR step to both bottleneck and copy our randomized linear DNA library.

Our CHANGE-seq-R approach has three key advantages compared to earlier approaches described by Pattanayak et al.[5]: 1) it is performed on circularized, partially randomized target libraries such that both parts of the cut and linearized target sequence can be readily sequenced after cleavage, 2) its read out is dependent on a single (rather than multiple) Cas9 cleavage events and 3) a controlled library size enables quantitative measurements of the enrichment of specific off-target sequences between control and nuclease treatments.

**Number of Unique Molecules.** After adding UMIs to the primers it was important to determine how many unique molecules could be surveyed that would allow adequate sequence depth of at least 10x coverage and ensure no collision issues, i.e. one UMI assigned to multiple off-target sequences. A 15bp UMI with an NNWNNW pattern, to reduce GC content, allows for approximately 33 million unique combinations. We determined that 3-4 million unique molecules limited collision effects and allowed an achievable sequencing target of approximately 50 million reads per sample.

#### Supplementary protocol 2: CHANGE-seq R

**SUPPLIES**

- IDTE pH 8.0 (1X TE Solution) (Integrated DNA Technologies, 11050204)
- Plasmid DNA p2T-CAG-eGFP-BlastR (modified from Addgene #107190)
- MilliporeSigma custom oligonucleotides
- Synthego custom sgRNA
- Proteinase K (NEB,P8107S)
- KAPA HiFi HotStart + ReadyMix (250 x 50 µl reactions) (Roche, KK2602)
- KAPA HiFi HotStart Uracil+ ReadyMix (250 x 50 µl reactions) (Roche, KK2802)
- DpnI (NEB R0176L)
- Ultrapure Nuclease Free water (Invitrogen 10977023)
- Magnum FLX magnetic rack (Alpaqua, A00400)
- T4 Polynucleotide Kinase (PNK) (NEB, M0201L)
- T4 DNA Ligase (NEB, M0202L)
- 10X T4 DNA ligase Buffer (NEB), supplied with T4 DNA Ligase
- USER Enzyme (NEB, M5505L)
- Exonuclease I (*E. coli*) (NEB, M0293L)
- Lambda Exonuclease (NEB, M0262L)
- Exonuclease III (E. coli) (NEB, M0206L)
- Plasmid-Safe ATP-dependent DNase (Epicentre, E3110K)
- 10X Ampligase Buffer (Lucigen, A1905B)
- 25mM ATP solution (Epicentre), supplied with Plasmid-Safe ATP-dependent DNase
- SrfI (NEB, R0629S)
- Quick CIP (NEB, M0525L)
- rCutsmart Reaction buffer (NEB, B6004SVIAL, supplied with Quick CIP and SrfI)
- SpCas9 (NEB, M0386T)
- Klenow Fragment (3' -> 5' exo-) (NEB, M0212L)
- NEBuffer 2 (NEB, B7002SVIAL supplied with Klenow Fragment)
- dNTPs (NEB, cat.no. N0447L)
- NEBNext® Ultra™ II Ligation Module (NEB E7595L)
- NEBNext® dA-Tailing Module (NEB E6053L)
- Kapa PEG/NaCl SPRI solution (Roche, 7961928001)
- NEBNext® Multiplex Oligos for Illumina® (Dual Index Primers Set 1) (New England BioLabs, E7600S)
- NEBNext adapter for Illumina (NEB), supplied with NEBNext® Multiplex Oligos for Illumina®
- Qubit dsDNA HS Assay Kit (Thermo Fisher Scientific, Q32854)
- NextSeq2000® kit 200-cycles (Illumina)
- Flow Cell, supplied with NextSeq2000® Reagent Kit
- NextSeq2000 RSB Buffer, supplied NextSeq2000® Reagent Kit
- PhiX Control V3 KIT (Illumina, FC-110-3001)
- Ultra pure EDTA 0.5 M (Invitrogen, 15575038)
- Ethanol (Sigma, cat.no. E7023)
- Sera-Mag Magnetic Beads; Carboxyl, Speedbeads; hydrophobic; 5 solids (Fisher/GE,.9981123)
- Guanidine thiocyanate (Sigma, G9277)
- Sodium Chloride 5 M Sterile (Invitrogen, AM9760G)
- TRIS Buffer 1.0 M solution, pH 8.0 (Fisher, 50146868)
- Polyethylene Glycol 8000 (Fisher, 507516674)
- Kappa Library Quantification Kit (Illumina/Uni) (Roche, KK4824)
- 1M Magnesium chloride (Invitrogen, AM9530G)
- Tween-20 (Sigma, P7949-500ML)
- HEPES (Fisher, BP310-1)

- Sodium Hydroxide solution (Sigma, 72068-100ML)
- 12well, 2% Agarose gel Cassettes (Yourgene Health, CG-10600-13-200(16))
- Ranger MQ Dual Dye Loading Buffer, 300 bp + 1k bp Marker (Yourgene Health, CG-14000-12-21)

**REAGENT SETUP**

**Resuspend the CHANGE-seq Randomized custom oligonucleotides.** For PCR1, forward mixed base primers were designed with a universal primer sequence, 15 base-pair unique molecular index (UMI) and partially randomized target sites to produce amplicons with a range of alterations at protospacer and PAM positions for each of the six target sites. For PCR2, Forward and Reverse ACG/deoxyUridine primers were designed to introduce a SrfI restriction site and ACGU sequences. Oligonucleotides were resuspended to 100 µM in TE pH 8.0. Resuspended oligonucleotides were kept at -20°C.

N = 25,25,25,25(GACT)
W=50.50(AT)
N1 = ACT (10%each) G=70%
N2 = GCT (10%each) A=70%
N3 = AGT (10%each) C=70%
N4 = ACG (10%each) T=70%

**oAF01_CTLA4.s9_FWD**
ACGCGAGCTGCATGTGTCAGANNWNNWNNWNNWNNW(N1)(N1)(N2)(N3)(N4)(N1)(N2)(N1)(N1)(N1)(N3)(N3)(N2)(N4)(N1)(N1)(N2)(N3)(N2)(N3)(N1)(N1)(N1)cttcttcaagtccgccatgc
**oAF02_AAVS1.s14_FWD**
ACGCGAGCTGCATGTGTCAGANNWNNWNNWNNWNNW(N1)(N1)(N1)(N1)(N3)(N3)(N2)(N3)(N4)(N2)(N1)(N1)(N1)(N2)(N3)(N2)(N1)(N1)(N2)(N4)(N4)(N1)(N1)cttcttcaagtccgccatgc
**oAF04_LAG3.s9_FWD**
ACGCGAGCTGCATGTGTCAGANNWNNWNNWNNWNNW(N1)(N2)(N2)(N1)(N1)(N3)(N4)(N1)(N2)(N1)(N2)(N4)(N3)(N3)(N4)(N1)(N1)(N2)(NI)(N1)(N1)(N1)(NI)cttcttcaagtccgccatgc
**oAF05_TRAC.s1_FWD**
ACGCGAGCTGCATGTGTCAGANNWNNWNNWNNWNNW(N1)(N4)(N3)(N2)(N1)(N1)(N1)(N4)(N4)(N3)(N4)(N1)(N1)(N2)(N4)(N2)(N4)(N3)(N4)(N1)(N4)(N1)(N1)cttcttcaagtccgccatgc
**oAF06_CCR5.s8_FWD**
ACGCGAGCTGCATGTGTCAGANNWNNWNNWNNWNNW(N1)(NI)(N2)(N3)(N2)(N1)(N4)(N2)(N2)(N1)(N2)(N2)(N1)(N1)(N2)(N2)(N2)(N2)(N2)(N3)(N2)(N1)(NI)cttcttcaagtccgccatgc

**o.AF07_PCR1_REV-** tgtgatcgcgcttctcgtt
**o.AF08_PCR2_FWD -** ACG/deoxyUridine/acgcgagctgcatgtgtcaga
**oAF09_PCR2_REV-** ACG/deoxyUridine/GCCCGGGCtctcgttggggtctttgctc

**Resuspend 1.5nmol gRNA from Synthego –** Resuspend in 15ul 1x TE buffer (10mM Tris, 1mM EDTA pH8.0 supplied with guides from Synthego). Aliquot and freeze at -80C. Avoid freeze/thaw cycles.

**CTLA4.s9** - GGACTGAGGGCCATGGACACGGG
**AAVS1.s14** – GGGGCCACTAGGGACAGGATTGG
**LAG3.s9** – GAAGGCTGAGATCCTGGAGGGGG
**TRAC.s1** – GTCAGGGTTCTGGATATCTGTGG
**CCR5.s8** - GGACAGTAAGAAGGAAAAACAGG

**Prepare Plasmid DNA**
Prepare plasmid DNA p2T-CAG-eGFP-BlastR via miniprep and dilute to ~1ng/ul.

**SPRI-guanidine binding buffer** 4M guanidine thiocyanate, 40mM TRIS, 17.6mM EDTA, pH 8.0. TRIS 1M pH 8 and EDTA 0.5M pH 8 can be added to the 4M guanidine (after the guanidine is solubilized in water – add the proper volume for getting the right final concentration) and then the pH will be very close to 8. Bring the pH to 8 with HCl.

**Sera-Mag Magnetic Beads preparation** Add 1 ml of Sera-Mag Magnetic Beads (Fisher/GE) to a 1.5 ml Eppendorf tube. Place in a magnetic rack. Remove the liquid. Remove the tube from the rack. Add 1 ml of TE and homogenize. Place back in the magnetic rack and remove the liquid. Repeat this step for a total of two TE pH 8.0 washes. Then, add 1 ml of TE pH 8.0. Note: this beads preparation step is required for preparing SPRI-guanidine beads and SPRI-beads.

**SPRI-guanidine beads preparation** Add 10 ml of 5M NaCl to 9 g of PEG 8000 and then add SPRI-guanidine binding buffer (prepared as described above) up to 49 ml. Homogenize for 5 min. Add 1 ml of Sera-Mag Magnetic Beads in TE (prepared as described above) and homogenize. Keep at 4°C.

**SPRI-beads preparation** Add 10 ml of 5M NaCl, 500 µl of 1M TRIS and 100 µl of 0.5M EDTA to 9 g of PEG 8000. Complete the volume to 49 ml with ultra-pure water. Add 1 ml of Sera-Mag Magnetic Beads in TE (prepared as described above) and homogenize. Add 27.5 µl of Tween-20 and homogenize. Keep at 4°C.

**10X Cas9 Nuclease Reaction Buffer –** 200mM HEPES, 1M NaCl, 10mM MgCl2,1mM EDTA; bring to pH 6.5 at 25°C with NaOH.

**PROCEDURE**

**PCR1-Creating highly diverse, partially randomized library for each target site**

**1|** Perform PCR1 for each target using the following components and PCR conditions.

| Component | Volume (µl) |
|---|---|
| 2x Kapa HiFi HotStart +Ready Mix | 25 |
| 10uM Mixed Base+eGFP_Fwd Primer | 1.5 |
| 10uM **o.AF07** PCR1 Rev Primer | 1.5 |
| p2T-CAG-eGFP-BlastR plasmid (1ng) | x |
| Nuclease-free water | x |
| Total | 50 |

| Step | Temperature | Time | Cycles |
|---|---|---|---|
| Denaturation | 95 °C | 3min | 1 |
| Denaturation | 98 °C | 20 s | 20 |
| Annealing | 73 °C | 15 s | 20 |
| Extension | 73°C | 30 s | 20 |
| Extension | 72 °C | 3min | 1 |
| Hold | 4°C | | 1 |

**2|** Add 1ul DpnI directly to PCR reaction. Incubate at 37°C for 15 minutes. Hold at 10°C

**3|** Dilute proteinase K 1:1 in water (2.5 µl of proteinase K and 2.5 µl of water) and add 5 µl of the dilution to PCR reaction. Incubate at 55 °C for 15 minutes.

**4|** Add 1.8X volumes (90 µl) of SPRI-guanidine beads to the PCR reaction, mix thoroughly by pipetting 10 times. Incubate at room temperature for 10 minutes. Place the reaction plate onto a Magnum FLX magnetic rack for 5 minutes. Remove the cleared solution from the reaction plate and discard. Add 200 µl of 80% ethanol, incubate for 30 seconds and remove the supernatant. Repeat this step for a total of two ethanol washes. Remove ethanol completely and let the samples air dry for 3 minutes on the magnetic rack. Remove the plate from the magnetic rack and add 40 µl of TE pH 8.0, and pipette 10 times to mix. Incubate at room temperature for 2 minutes. Place the reaction plate back to the magnetic rack for 1 minute. Transfer the eluted DNA to a new plate.

*Note:* using SPRI-Guanidine beads in this purification step to completely inactivate and remove Kapa HiFi HotStart + polymerase will help increase yield of circularized DNA. Carryover of this enzyme will remove the 3' overhangs generated in step 9, due to its strong 3'-5' exonuclease activity.

**5|** Quantify PCR products with Qubit. Dilute each sample to a concentration of 1pg/ul.

***Safe Stop – store purified PCR products at 4°C or -20°C***

**PCR2-Bottlenecking and making copies of library**

**6|** Perform PCR2 with 1pg/ul dilutions of PCR1 products for each target using the following components and PCR conditions. **Use Uracil tolerant polymerase** to synthesize ACGU sequences.

| **Component** | **Volume (µl)** |
|---|---|
| 2x Kapa HiFi HotStart **Uracil** +Ready Mix | 25 |
| 10uM **o.AF08** PCR2 Fwd Primer | 1.5 |
| 10uM **o.AF09** PCR2 Rev Primer | 1.5 |
| **PCR1 product (1pg)** | x |
| Nuclease-free water | x |
| Total | 50 |

| **Step** | **Temperature** | **Time** | **Cycles** |
|---|---|---|---|
| Denaturation | 95 °C | 3min | 1 |
| Denaturation | 98 °C | 20 s | 30 |
| Annealing | 72 °C | 15 s | 30 |
| Extension | 72°C | 15 s | 30 |
| Extension | 72 °C | 3min | 1 |
| Hold | 4°C | | 1 |

**7|** Dilute proteinase K 1:1 in water (2.5 µl of proteinase K and 2.5 µl of water) and add 5 µl of the dilution to PCR reaction. Incubate at 55 °C for 15 minutes.

**8| Purify -** Add 1.8X volumes (90 µl) of SPRI-guanidine beads to the PCR reaction, and follow purification as described in **step 4**. Add 40 µl of TE pH 8.0 to elute and transfer the eluted DNA to a new plate.

**9|** USER/PNK. Set up the USER/PNK reaction as follows.

| **Component** | **Volume (µl)** |
|---|---|
| T4 DNA Ligase Buffer (10X) | 5 |
| USER Enzyme (1 U/µl) | 3 |
| T4 Polynucleotide Kinase (10 U/µl) | 2 |
| Purified DNA (no more than 8ug) | 40 |
| Total | 50 |

Incubate in thermocycler at 37°C for 1 hour.

**10| Purifty -** Add 1.8X volumes (90 µl) of SPRI beads to the PCR reaction, and follow purification as described in **step 4**. Add 35 µl of TE pH 8.0 to elute and transfer the eluted DNA to a new plate.

**11|** Run each sample on a QIAxcel capillary electrophoresis instrument, in a 0.2 ml thin-walled 12-well strip tube with a QIAxcel DNA High Resolution Kit (Qiagen), QX Alignment Marker 50 bp to 1.5kb (Qiagen) and QX Size Marker 15bp – 3kb (Qiagen), following manufacturer's instruction. The average size should of DNA should be ~466 bp.

**12|** Quantify by Qubit dsDNA High sensitivity assay. Calculate 500ng of DNA per reaction. Use TE pH8.0 to bring each reaction volume to 88ul. Save any remaining DNA not used for circularization to run as control on QIAxcel in step 19.

*Note:* Number of samples will likely double at this step.

**13|** Intramolecular circularization. Set up the DNA circularization as follows:

| **Component** | **Volume (µl)** |
|---|---|
| T4 DNA Ligase Buffer (10X) | 10 |
| T4 DNA Ligase (400 U/µl) | 2 |
| USER/PNK treated DNA (500 ng) | *variable* |
| IDTE pH 8.0 | *variable* |
| Total | 100 |

Incubate in thermocycler at 16°C for at least 16 hours (overnight).

**14| Purify** the circularized DNA reactions as previously described in step 4 by adding 1X volumes (100µl) of SPRI beads to the DNA. Elute in 37 µl of TE pH 8.0. Transfer eluted DNA to a clean plate.

**15|** Plasmid-Safe ATP-dependent DNase/Lambda Exo/ExoI/ExoIII treatment:

| **Component** | **Volume (µl)** |
|---|---|
| Ampligase Buffer (10X) | 5 |
| ATP (25 mM) | 2 |
| Plasmid-Safe ATP-Dependent DNase (10 U/µl) | 2 |
| Lambda Exonuclease (5 U/µl) | 2 |
| Exonuclease I (*E. coli*) (20 U/µl) | 1 |
| Exonuclease III (*E. coli*) | 1 |
| Circularized DNA | 37 |
| Total | 50 |

Incubate in a thermocycler at 37 °C for 1 h. Add 5 µl of EDTA 0.5 M, incubate at 70 °C for 30 min, hold at 4 °C.

**16| Purify** the circularized, exonuclease treated DNA reactions as previously described in step 4 by adding 1X volumes (50 µl) of SPRI beads to the DNA. Elute with 43 µl of TE pH 8.0. Transfer the supernatant to a new plate.

**17|** Quick CIP treatment of circularized exonuclease-treated DNA. Set up Quick CIP reaction as follows:

| **Component** | **Volume (µl)** |
|---|---|
| CutSmart Reaction Buffer (10X) | 5 |
| Quick CIP (5 U/µl) | 2 |
| Exonuclease Treated DNA | 43 |
| Total | 50 |

Incubate at 37 °C for 10 min. Heat inactivate at 80 °C for 2 min.

**18| Purify** CIP treated, circularized DNA as previously described in step 4 by adding 1.8X volumes (90 µl) of SPRI-beads to the DNA. Elute with 10µl of TE pH 8.0. Transfer the supernatant to a new plate, pool all reactions of the same sample type, and quantify by Qubit HS assay.

**19| QC Step:** Run each sample on QIAxcel with 6ul circularized DNA. Run saved USER/PNK treated DNA as a size control. Circularized DNA should run faster on the gel.

***Safe-Stop-Circularized, Exonuclease and CIP treated DNA can be stored at -20°C***

**Cleavage of enzymatically purified, circularized plasmid DNA**

**20|** sgRNA dilution and re-fold. Dilute the sgRNA to 9 µM in 10ul nuclease-free water and use the follow program on a thermocycler for sgRNA re-fold:

| **Step** | **Temperature** | **Time** | **Cycles** |
|---|---|---|---|
| 1 | 90 °C | 5 min | 1 |
| 2 | 90-25 °C | Ramp rate 2% | |
| Hold | 4°C | | 1 |

**21|** *In vitro* cleavage with S.py Cas9. Perform step **21** concurrently with step **22** (SrfI cleavage).

**A)** Dilute Cas9 to 1uM

| **Component** | **Volume (µl)** |
|---|---|
| Cas9 Nuclease Reaction Buffer (10X) | 6 |
| NEB S.py Cas9 (20uM) | 3 |
| Nuclease free H20 | 51 |
| Total | 60 |

**B)** Prepare RNP complex with Cas9 and sgRNA. Setup *in vitro* master-mix:

| **Component** | **Volume (µl)** |
|---|---|
| Cas9 Nuclease Reaction Buffer (10X) | 5 |
| S.py Cas9 dilution mix **(A)** (1uM) | 4.5 |
| sgRNA-folded (9 µM) | 1.5 |
| Total cleavage master-mix | 11 |

Incubate at room temperature for 10 min.

**C)** *In vitro* cleavage. Add circularized DNA (125 ng, volume varies) and nuclease-free water to complete the 50 µl:

| | |
|---|---|
| Cleavage master-mix | 11 |
| Exonuclease Treated DNA (125 ng) | *X* |
| Nuclease-free water | *X* |
| Total | 50 |

Incubate in a thermocycler at 37 °C for 1 h, hold at 4 °C.

**22|** Prepare reaction for control restriction enzyme SrfI cleavage.

**A)** Prepare Master mix.

| Component | Volume (µl) |
|---|---|
| Neb rCutsmart Buffer (10X) | 5 |
| NEB SrfI | 1 |
| NFW | 5 |
| Total cleavage master-mix | 11 |

Incubate at room temperature for 10 min.

**Important -** DNA must come from same pooled circularized sample as step **21** for each target. (125 ng, volume varies) and nuclease-free water to complete the 39 µl.

| Component | Volume (µl) |
|---|---|
| Cleavage Master-mix | 11 |
| Circularized DNA (125ng) | *X* |
| Nuclease-free water | X |
| Total volume | 50 |

Incubate in thermocycler at 37 °C for 40 minutes, then 65 °C for 20 minutes, hold at 4 °C.

**19|** Dilute proteinase K 1:5 in water (1 µl of proteinase K and 4 µl of water) and add 5 µl of the dilution to the *in vitro*-cleaved DNA and incubate in a thermocycler at 37 °C for 15 min.

**20| Purify** cleaved (Cas9 and SrfI) DNA as previously described in step 4 by adding 1X volumes (55 µl) of SPRI-beads to the DNA. Elute with 40µl of TE pH 8.0. Transfer the supernatant to a new plate.

**21| End repair**. Setup the end-repair master mix:

| Component | Volume (µl) |
|---|---|
| NEB Buffer 2 (10X) | 5 |
| Klenow fragment (3'->5' exo-) (5 U/µl) | 1 |
| dNTP | 4 |
| Total end-repair master-mix | 10 |

Add 10 µl end-repair master-mix to each eluted DNA sample.

| | |
|---|---|
| End-repair master-mix | 10 |
| cleaved DNA | 40 |
| Total | 50 |

Incubate on a thermocycler at 37 °C for 30 min, then 75 °C for 20 min, hold at 4 °C.

**22| Purify** end-repaired DNA as previously described in step 4 by adding 1X volumes (50 µl) of SPRI-beads to the DNA. Elute with 42µl of TE pH 8.0. Transfer the supernatant to a new plate. **Keep the beads.**

**23| A-tailing**. Setup the A-tailing master mix

| **Component** | **Volume (µl)** |
|---|---|
| NEBNext dA-Tailing Reaction Buffer | 5 |
| Klenow Fragment (3′ → 5′ exo–) | 3 |
| Total A-tailing master-mix | 8 |

Add 8 µl of A-tailing master-mix to each eluted DNA sample with beads.

| | |
|---|---|
| A-tailing master-mix | 8 |
| Cleaved DNA/beads | 42 |
| Total | 50 |

Incubate on a thermocycler at 37 °C for 30 min, hold at 4 °C.

**24| Purify** A-tailed DNA as previously described in step 4 by adding 1.8X volumes (90 µl) of SPRI-beads to the DNA. Elute with 60µl of TE pH 8.0. Transfer the supernatant to a new plate. **Keep the beads.**

**25|** Adapter ligation. Setup the adapter ligation master-mix

| **Component** | **Volume (µl)** |
|---|---|
| NEBNext Ultra II Ligation Master Mix | 30 |
| NEBNext Ligation Enhancer | 1 |
| Total MM | 31 |
| End Prep Reaction Mixture/beads | 60 |
| NEBNext Adapter for Illumina (15 µM) | 2.5 |
| Total reaction volume | 93.5 |

Incubate on a thermocycler at 20 °C for 15 minutes **with heated lid off,** hold at 4 °C.

*Note*: Prepare single-use aliquots of NEB adapters to avoid adapter dimer formation due to freeze–thaw hydrolysis of the 3′ T′

**26| Purify** Add 1X volumes (50 µl) of PEG/NaCl solution to the adapter-ligated DNA and purify DNA as described in step 4. Elute in 47 µl of TE pH 8.0 and **keep the beads.**

**27|** USER enzyme. Add 3 µl of USER enzyme, provided with NEBNext® Multiplex Oligos for Illumina® (Dual Index Primers Set 1) to the adapter ligated DNA with beads. Incubate at 37 °C for 15 min.

**28| Purify** Add 0.7X volumes (35 µl) of PEG/NaCl solution to the USER Enzyme treated DNA and purify as previously described in step 4. Elute in 20 µl of TE pH 8.0. Transfer the supernatant to a new semi-skirted PCR plate and quantify by Qubit dsDNA HS assay and proper Qubit assay tubes (usually about 1-5 ng/µl).

**29|** PCR. Prepare PCR master-mix for adding dual-index barcodes:

**A)**

| Component | Volume (µl) |
|---|---|
| 2x Kapa HiFi HotStart Ready Mix | 25 |
| NEBnext i5 Primer (10uM) | 5 |
| NEBnext i7 Primer (10uM) | 5 |
| **USER treated DNA (10-20ng)** | x |
| Nuclease-free water | x |
| Total | 50 |

*Note:* Carefully record which primers were used for each

**B)** Perform the PCR using the following thermocycling conditions:

| Step | Temperature | Time | Cycles |
|---|---|---|---|
| Denaturation | 98 °C | 45 s | 1 |
| Denaturation | 98 °C | 15 s | 20 |
| Annealing | 65 °C | 30 s | 20 |
| Extension | 72 °C | 30 s | 20 |
| Extension | 72 °C | 1 min | 1 |
| Hold | 4°C | | 1 |

***Safe-Stop. PCR can hold at 4°C overnight ***

**30| Purification**. Add 0.7X volumes of SPRI-beads to the PCR and purify as previously described in step 4. Elute in 30 µl of TE pH 8.0. Transfer the supernatant to a new semi-skirted plate.

*Note:* Can run samples on Qiaxcel to determine purity and size of NGS library prepared samples.

**31| Quantify library for sequencing.** Make 1:10 serial dilutions of 20 µl from $10^{-1}$ to $10^{-7}$ dilution of each sample from the library (PCR), starting with 2µl of DNA and 18 µl of TE, and mix well.

**A)** Assemble qPCR with Kapa library quantification kit. Prepare master-mix solution as follows:

| Component | 1 reaction (µl) | Final Concentration |
|---|---|---|
| KAPA SYBR FAST qPCR Master Mix (2X) + Primer Premix (10X) | 12 | 1X |
| Nuclease-free water | 4 | |
| Total qPCR mix | 16 | |

**B)** Assay 4 different dilution factors (4 µl) for each sample ($10^{-4}$, to $10^{-7}$ from the library) in duplicate (in an appropriate 96-well plate). A standard curve (provided with Kapa Library Quantification Kit) and a non-template control (NTC) are required. Add 4 µl of each standard in duplicate, and nuclease-free water in the NTC. Add 16 µl of qPCR master-mix to each sample.

| Component | Volume (µl) | Final concentration |
|---|---|---|
| qPCR mix | 16 | |
| Sample (add nuclease-free water into the NTC well) | 4 | *variable* |
| Total | 20 | |

**C)** Seal the plate and spin down. Run qPCR in appropriate thermocycler with the following program:

| Cycling step | Temperature | Time | Cycles |
|---|---|---|---|
| Initial denaturation | 95 °C | 5 min | 1 |
| Denaturation | 95 °C | 30 s | 35 |
| Annealing/extension/data acquisition | 60 °C | 45 sec | 35 |
| Melt curve analysis | 60-95 °C | | |

**D)** Add the appropriate DNA copies for each standard when setting up the qPCR plate in the qPCR program, as follows:

| Standard | dsDNA molecules/µl |
|---|---|
| Standard 1 | $1.2 \times 10^7$ |
| Standard 2 | $1.2 \times 10^6$ |
| Standard 3 | $1.2 \times 10^5$ |
| Standard 4 | $1.2 \times 10^4$ |
| Standard 5 | $1.2 \times 10^3$ |
| Standard 6 | $1.2 \times 10^2$ |

**32|** Analyze qPCR results. Multiply the average of duplicate values by the dilution factor and by the five-fold dilution factor of the qPCR reaction, as follows: Total copies/µl = # * dilution factor.

**33|** Pool library for NextSeq2000. Pool all the samples in one library at equimolar concentrations. 1X pooled library should be in a total volume of 5 µl, ~ $8 \times 10^9$ molecules.

**34| Size selection –** to remove smaller DNA fragments, use 2% agarose gels and 300-1kb marker from Yourgene Health. Size select 500~900bp fragment of pooled library with LightBench automated size selection instrument.

**35|** Confirm size of library by running pooled, sized selected sample on Qiaxcel**.**

**36|** Check concentration with Qubit High Sensitivity Assay. Use the size of the library calculated from step 35. Concentration should be around 2nM.

**37| Loading the sample for sequencing.**

**A)** Dilute library to 625pM with RSB buffer (supplied with Nextseq2000 kits) for a total of 24ul.
**B)** Prepare the Phix control V3 (PhiX Control V3 KIT) by mixing 2ul of 10nM PhiX control with 8ul RSB buffer for a total of 10ul
**C)** Take 7.8 ul of PhiX mix from step **B** and mix with 16.2ul RSB for a total of 24ul
**D)** Take 20.4ul of library mix from step **A** and mix with 3.6ul PhiX mix from step **C.**

**35|** Load and sequence library using a NextSeq2000 200-cycles kit according to manufacturer's instructions using NextSeq 2000 system. Sequencing is performed with 100 bp paired-end reads and 8 bp dual-index reads. At least 10x coverage of each sample is required.

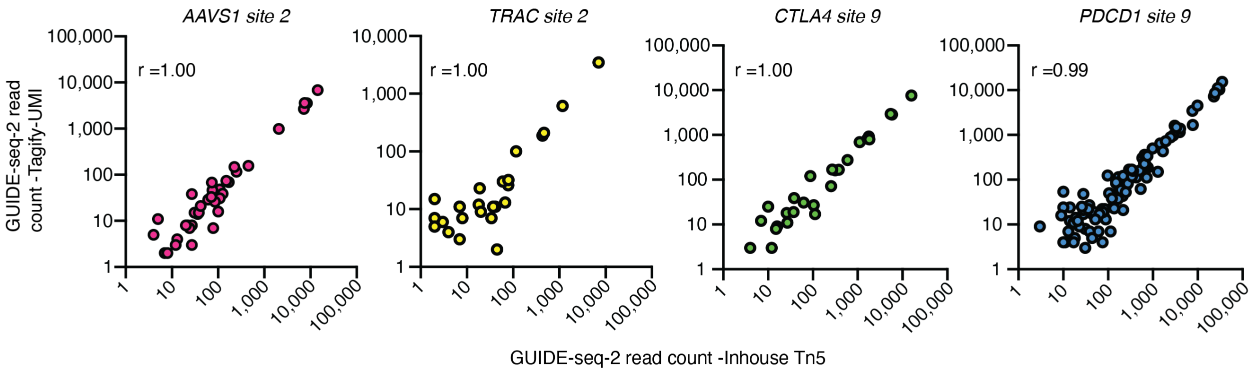

**Supplementary figure 1| GUIDE-seq-2 profile correlates well between inhouse Tn5 and commercially available Tn5** Scatterplots of GUIDE-seq-2 read counts (log10) from experiments performed on primary human T cells for 4 target sites. Genomic DNA were tagmented using inhouse Tn5 (x-axis) and seqWell Tagify-UMI (y-axis) before GUIDE-seq-2 PCR. GUIDE-seq-2 library prepared with each method was sequenced individually with NextSeq2000. r = Pearson's correlation coefficient.

St. Jude - Confidential

#### Supplementary Tables

**Supplementary Table 1. gRNAs**
*See attached file.*

**Supplementary Table 2. Coriell cell lines**
*See attached file.*

**Supplementary Table 3. rhAmp-seq panels**
*See attached file.*

**Supplementary Table 4. GUIDE-Seq-2 data**
*See attached file.*

**Supplementary Table 5. CHANGE-Seq-R Primers**
*See attached file.*

**Supplementary Table 6. CHANGE-net 1000 Genomes predictions**
*See attached file.*

St. Jude - Confidential
