## Supplementary material for "Frequent, context-dependent effects of human genetic variation on Cas9 activity revealed by population-scale GUIDE-seq-2 and deep combinatorial CHANCE-seq profiling": Online Methods

**Cell culture.** Lymphoblastoid cells (LCLs) from 94 donors across four genetic backgrounds were purchased from Coriell Institute (**Supp. Table 2**). LCLs were individually cultured in RPMI-16 (Thermo Fisher Scientific), supplemented with 10% FBS and penicillin-streptomycin (50 U ml^-1^) (Thermo Fisher Scientific) at 37 °C with 5% CO_2._ Human primary CD4^+^/CD8^+^ T cells were cultured in X-Vivo 15 medium (Lonza) supplemented with 5% human heat-inactivated serum (HSA) (Fisher), 10 ng ml^−1^ IL-7 (Miltenyi) and 10 ng ml^−1^ IL-15 (Miltenyi). T cells were stimulated with MACS GMP T cell TransAct polymeric nanomatrix (Miltenyi) for 3 days according to the manufacturer’s instructions before transfection.

**Cell transfection.** Transfection of lymphoblastoid cell lines were performed with Cas9-sgRNA ribonucleoprotein (**RNP**) complex containing 20 pmol of purified recombinant Cas9 (3x-NLS) (Protein Production Core Facility, St. Jude) and 9-fold molar excess of synthetic chemically modified sgRNA (Synthego), in SF solution (Lonza). 100 pmol of end-protected dsODNs were added directly to the cell suspension before nucleofection. RNPs were added directly to 2.5 x 10^5^ cells resuspended in 20 μl of SF solution and nucleofected with preprogrammed pulse DN-100 in a 4D-Nucleofector system (96-well unit) (Lonza). After nucleofection, cells were recovered in RPMI-16 medium with 20% FBS. Cell population was assessed by NucleoCounter NC-3000 (ChemoMetec) with acridine orange (AO) and 4’,6-diamidino-2-phenylindole (DAPI) AO-DAPI staining reagent 3 days post nucleofection. After 5 days, cells were collected for genomic DNA purification.

Human primary CD4^+^/CD8^+^ T cells were nucleofected with Cas9–sgRNA RNP complex containing 75 pmol of purified recombinant Cas9 (Protein Production Core Facility, St. Jude) and 3-fold molar excess of synthetic sgRNA (Synthego). RNPs were added directly to 6 × 10^5^ cells resuspended in 20 μl of P3 solution containing 100 pmol of end-protected dsODNs (for GUIDE-seq-2), and nucleofected with programmed pulse code EO-115 in a 4D-Nucleofector System (Lonza). After nucleofection, cells were recovered in X-Vivo 15 medium with 20% human heat-inactivated serum (HSA) (Fisher), 10 ng ml^−1^ IL-7 (Miltenyi) and 10 ng ml^−1^ IL-15 (Miltenyi). After 3 days, cells were collected for genomic DNA purification.

**GUIDE-seq-2 high-throughput cellular profiling.** Genomic DNA from edited and wild-type control LCLs cells or T cells containing the dsODN tag was isolated using DNAdvance Kit (Beckman Coulter) according to manufacturer's instructions. Purified genomic DNA was quantified by Quant-iT BR and 100 ng of gDNA was tagmented with a custom Tn5-transposome containing a 10-nt UMI to an average length of 600 bp and purified using SPRI-guanidine magnetic beads. Tagmented gDNA was PCR-amplified using dsODN sense and antisense-specific primers in separate reactions for each sample for target enrichment and then purified with a limited amount of SPRI magnetic beads, for one-step purification and normalization. Libraries for the same sgRNA target site or controls were pooled and quantified by qPCR (KAPA Biosystems) and sequenced with 146-bp paired-end reads on an Illumina NextSeq or NovaSeq with approximately 15-20% PhiX to account for low-diversity library sequencing. GUIDE-seq-2 library preparation protocol was adapted to an automated liquid-handling workflow (Biomek) to process the over 600 samples generated in this study. A detailed user protocol for GUIDE-seq-2 is provided (**Supplementary Protocol 1**).

GUIDE-seq-2 data analyses were performed using open-source GUIDE-seq-2 analysis software (<https://github.com/tsailabSJ/guideseq>, v2.0.0). The parameters were: demultiplex mismatch: 1; window_size: 25; mapq_threshold: 50; max_score: 7. The GUIDE-seq-2 pipeline first demultiplexes the samples using sample barcode (allowing 1 mismatch), extracts UMIs , and then aligns the reads to the genome using BWA mem with default parameters. SAMtools is used to filter out reads based on MAPQ threshold. UMItools is used to remove PCR duplicates. Off-targets were identified using identifyOfftargetSites.py. Specifically, peaks of GUIDE-seq-2 read counts were called and extended upstream and downstream for 25bp. Off-target sites were identified using the sequence homology search between the above window sequence and the on-target sequence. A distance score of <=7 was required as well as at least two GUIDE-seq-2 read counts, where within one sample these reads map bidirectionally from the DNA double-stranded break point, or where they map in the same direction but are amplified from two different primers. This stringent requirement for two molecularly distinct events within a small genomic window minimizes false positives. The distance score is a weighted score based on the number of mismatches and the number of indels, where one indel is counted as 3.

**Multiplex targeted amplicon sequencing.** To determine the indel mutation frequency at GUIDE-seq-2 identified sites harboring SNPs, off-target sites were amplified from genomic DNA from edited cells or unedited controls using the rhAmpSeq system (IDT), in triplicates, with primers listed in **Supplementary Table 3**. Sequencing libraries were generated according to manufacturer’s instructions. Completed libraries were quantified by qPCR using Kapa Library Quantification kit (Kapa Biosystems) and sequenced with 151-bp paired-end reads on an Illumina MiSeq or NextSeq instrument. We originally submitted 304 on- and off-target sites to IDT for panel design. The final multiplexed targeted sequencing panel (rhAmpSeq) included 284 sites, reflecting an approximate 7% dropout rate, due to failure to amplify certain genomic regions. During downstream data analysis, we required a minimum of 1,000 reads per site per donor, which resulted in 283 sites passing quality control.

**Indel and targeted tag sequencing analysis.** Paired-end high-throughput sequencing reads were processed to remove adapter sequences with trimmomatic^1^ (version 0.36), merged into a single read with FLASH^2^ (version 1.2.11) and mapped to human genome reference hg38 using BWA-MEM^3^ (version 0.7.12). Reads that mapped to on- or off-target sites were realigned to the intended amplicon region and to the dsODN tag sequence using a striped Smith–Waterman algorithm as implemented in the Python library scikit-bio; indels and tag integrations were counted and reported with total read counts. Code was available on <https://github.com/tsailabSJ/count_indels_integrations>.

**Genetic data analysis.** High-coverage variants variant call files (VCF) files were downloaded from the 1000 genomes project^4^ and the GNOMAD v4.1 database. Variants-containing off-targets for G1000 VCF files were obtained using PLINK (v2.0) with “--extract bed0” option and for GNOMAD VCF files were obtained using bcftools (v1.17). GUIDE-seq-2 read counts were normalized as counts per million based on the sum of on and off-targets for each sample. PLINK (v2.0) was applied to perform association analysis (--glm) for each variant with GUIDE-seq-2 normalized read count (log2-transformed), rhAmpSeq indel frequency, and rhAmpSeq dsODN integration frequency. FDR was calculated using the BH method. A high-confidence variant dataset was generated as follows. The genotype association (GA) test scores were set as 1 if their FDR<=0.1 and the absolution variant effect (beta score from PLINK) >= 0.5; otherwise 0. The allelic imbalance scores (a measure of whether one allele was preferentially edited) were calculated based on rhAmpSeq data for all heterozygous donors. For rhAmpSeq indel data, allelic imbalance was calculated only using reads containing indels. Fisher exact test was applied based on 2 by 2 contingency table: row 1: read counts for the reference allele, indel reads and non-indel reads. Row 2: read counts for the alternative allele, indel reads and non-indel reads. For rhAmpSeq dsODN data, AI score was calculated similarly but with only reads containing the dsODN primer. FDR was calculated using the BH method. The AI scores were set as 1 if their FDR <=0.1. The high-confidence dataset is defined as a variant having total GA scores >=2, or GA>=1 & AI>=1. For high-confidence variant, the minimal number of donors harboring the variant allele (i.e., heterozygous donors plus homozygous alternative donors) is 2. There are 45 variants with no rhAmp-Seq data, therefore not evaluated for high-confidence.

**Analysis of GUIDE-seq-2 sensitivity.** GM24631 LCLs were edited with Cas9 or HiFi-Cas9 targeting the AAVS1 locus (target sequence: GTCCCTAGTGGCCCCACTGT) in the presence of the GUIDE-seq-2 dsODN tag as described elsewhere. Single cell clones were established to isolate ‘off-target clones’ with both on- and off-target tag integration and ‘on-target clones’ with only on-target tag integration from the Cas9 and HiFi-Cas9 edited cells, respectively. gDNA from each clone were diluted to 5 ng/uL with IDTE and the gDNA of an ‘off-target clone’ was serially diluted with the gDNA of an ‘on-target clone’. 100 ng of each mixture were subjected to GUIDE-seq-2 to compare the expected fraction of off-targets and GUIDE-seq-2 readcounts. Lower limit of detection (LLOD) was determined where GUIDE-seq-2 cannot detect one or more of the replicates (n=3) of the off-targets. Lower limit of quantification (LLOQ) was determined as the lowest fraction of off-targets where GUIDE-seq-2 readcounts were reproducible from all the replicates.

**CHANCE-seq massively parallel biochemical profiling.** Plasmid DNA (Addgene #107190) that was modified to p2T-CAG-eGFP-BlastR with Gibson Assembly was arbitrarily chosen as DNA template for PCR1. PCR1 was performed with KAPA Hifi Hotstart + Ready Mix(Roche), plasmid DNA, and forward mixed base primers (MilliporeSigma) containing a universal primer sequence, a 15 base pair unique molecular index (UMI), and a partially randomized target site with a 30% chance of mismatch at each of the 23 bases across protospacer and PAM positions for each of five target sites (AAVS1 site 14, *CTLA4* site 9*, CCR5* site 8, *LAG3* site 9 and *TRAC* site 1) and a reverse primer that was complimentary to the plasmid backbone to make a 466bp amplicon. After PCR1, the reactions were treated with *DpnI* (NEB) and proteinase K (NEB), then bead-purified. Concentrations for all PCR products were measured via Qubit High sensitivity assay (Thermo Fisher) and diluted to 1pg/ul. The library was bottlenecked to approximately 3 million molecules or 1pg of DNA for use in PCR2. PCR2 was performed with KAPA Hifi Hotstart Uracil + Ready Mix (Roche), a forward primer that introduces an ACGU sequence, and a reverse primer that introduces an ACGU sequence and a *SrfI* restriction site. After PCR2, the reactions were treated with proteinase K (NEB), bead-purified then treated with USER enzyme (NEB) and T4 PNK (NEB). Intramolecular circularization was performed by combining bead-purified DNA with T4 DNA ligase (NEB) in T4 DNA ligase buffer. Circularized DNA was purified and then treated with a cocktail of exonucleases containing Exonuclease I (NEB), Lambda Exonuclease (NEB), Exonuclease III (NEB) and Plasmid-safe DNase (Lucigen). Quick CIP (NEB) was used on exonuclease treated DNA. *In vitro* cleavage reactions were performed with 125 ng of exonuclease, CIP-treated, circularized DNA, 90 nM of SpCas9 protein (NEB) and 270 nM of sgRNA, in a 50 μL volume with 1x Cas9 reaction buffer (20 mM HEPES; 100 mM NaCl; 1 mM MgCl2; 0.1 mM EDTA; pH 6.5 @ 25°C.) Control samples were cleaved with *SrfI*. Cleaved products in Cas9 and control groups were end-repaired, A-tailed, ligated with a hairpin adaptor (NEB), treated with USER enzyme (NEB) and amplified by PCR with dual-index NEBNext Multiplex Oligos for Illumina (NEB), using KAPA HiFi Hotstart + Ready Mix (Roche). Libraries were quantified by qPCR with KAPA library quantification kit (Roche), pooled, size selected, and sequenced with 100 bp paired-end reads on an Illumina NextSeq2000® instrument. A detailed user protocol for CHANCE-seq is provided (**Supplementary Protocol 2**) along with **Supplementary Note 2** describing key optimization steps.

**CHANCE-Seq Data Processing and Analysis**. Sequence data analyses were performed using open-source CHANCE-seq analysis software (https://github.com/tsailabSJ/CHANCEseq). FASTQ reads of cleaved products were processed using SeqKit (version 2.5.1) to stitch paired-end reads^5,6^. During this process, the R1 reads were reverse-complemented to ensure correct orientation. The 15 bp unique molecular identifier (UMI) and the subsequent 23 bp target sequence were extracted using Cutadapt (version 4.4), guided by specific 10 bp flanking sequences: ATGTGTCAGA for the 5’ end and CTTCTTCAAG for the 3’ end. Both forward and reverse complement sequences were considered during the search. An error tolerance of 0.1 was applied, requiring at least 9 out of the 10 bp of the flanking sequences to match. Targets were retained if their lengths were 19–27 bp (23 ± 4) to accommodate end-repair products in Cas9-treated samples.

To correct sequencing errors and deduplicate UMIs, singleton reads (reads observed exactly once) were reassigned to a unique 1-edit-distance neighbor only when exactly one such higher-count neighbor exists; otherwise, the singleton was left unchanged. In control samples, barcodes mapping to multiple randomized targets were collapsed to the dominant sequence (highest read count) provided each alternative was within an edit distance ≤2 of that dominant sequence. This two-step procedure corrects (i) sequencing errors leading to UMI duplication and (ii) minor sequence errors in the target linked to a barcode, while minimizing unintended merges across genuinely distinct molecules. Additionally, to ensure high-quality data, the corrected UMIs were required to match the pattern NNWNNWNNWNNWNNW, where N represents any nucleotide and W represents A or T. Then, using corrected control data, we identified collision-free barcodes, defined as barcodes associated with only one unique randomized target of 23 bp in length and stored them in a dictionary.

In Cas9-treated samples, targets were required to be within an edit distance of 3 or less from the corresponding target in the control samples for the same barcode. This threshold further minimizes the risk of barcode collision, where two different targets share the same barcode due to random chance in randomized target library synthesis. An edit distance of 3 or less allows for variations such as one sequencing error and up to two end-repair insertions, allowing for a balance between error tolerance and specificity. These barcodes were then used to evaluate the presence of cleaved molecules in Cas9-treated samples. Reads that did not meet these quality control criteria were discarded. Unique targets were counted based on their associated barcodes, and multiple barcodes mapping to the same target were combined.

The reads were normalized to counts per million (CPM). Log-transformed fold changes were calculated as log2(1 + FC), where FC represents the fold change of Cas9 CPM relative to control CPM. This transformation helps stabilize variance and accommodates small fold change values without perturbing rank order. Percent relative activity is defined as (fold change off-target/fold change on-target)*100, where fold change is defined as above. Randomized targets with less than five raw control reads and more than 6 mismatches were excluded from our analysis.

For the PAM specificity heatmap, only randomized targets with mismatches in the PAM and no mismatches in the protospacer were considered. Average relative activity for both replicates was calculated for each target separately. The number of possible sequence combinations for each mismatch count was determined using the equation $(22/n)*3^{2}*4$, where $n$ is the number of mismatches, 22 is the number of bases that could have a mismatch, 3 is the number of alternate base, and 4 is for the N in NGG where it’s not considered as a mismatch, but still counted towards the number of unique combinations. For quantifying mismatch effects by position, Cas9 reads were normalized to control counts and standardized to 100% within each mismatch group (mismatch levels 1-6) and the average relative activity are plotted. In the global analysis of CHANCE-Seq mismatches, relative activity was finely binned, and the normalized number of mismatches (transversions or transitions) at each position (excluding position 21) was displayed with up to 50% opacity to distinguish each type. For sequence visualizations of high and low relative activity, the top 10 sequences with the highest relative activity were shown. If more than 10 sequences tied for the lowest activity of 0, sequences were randomly sampled from this group.

**Mismatch Synergy Calculations**

The synergistic effect between two mismatches was quantified in two steps. First, the individual effect of each single mismatch was estimated as the ratio of the experimentally measured relative activity of the off-target sequence containing that mismatch to the on-target activity. Second, for each pair of mismatches, the expected activity under the assumption of independence was calculated by multiplying the individual effects of the two single mismatches. The observed activity was defined as the ratio of the experimentally measured activity of the off-target sequence containing both mismatches to the on-target activity. The synergistic effect was then quantified as the log2 ratio between the expected and observed activity changes.

**Deep learning model**

To prepare the CHANCE-seq data for model training, the following preprocessing steps were applied. First, relative activity was calculated as the ratio of off-target fold change to on-target fold change. This ratio was then log-transformed using the formula: log2(1 + relative activity). Experimental replicates were combined by averaging their log-transformed relative activity. To reduce noise, off-target sequences with control read counts below 5 were excluded from model training. A high-confidence test set was created by randomly selecting 10% of the off-target sequences with at least 10 control read counts. For the remaining training dataset, an additional 10% was further set aside for validation to optimize hyperparameters and prevent overfitting. Input features were generated by concatenating one-hot encoded on-target and off-target sequences, each 23 bp in length, resulting in an input matrix with dimensions (23, 8). This matrix served as the input for the deep learning model.

A 6-layer Convolutional Neural Network (CNN) was constructed to predict the log-transformed fold change in CHANCE-seq data. The model architecture consists of five convolutional layers, each with 128 kernels, followed by a final convolutional layer with 10 kernels. Kernel sizes for the convolutional layers are set to (3, 3, 3, 5, 5, 5), respectively. Batch normalization is applied after each convolutional layer, and LeakyReLU is used as the activation function. Dropout with a rate of 0.2 is applied to prevent overfitting. The output from the convolutional layer is flattened and passed through two fully connected layers with 128 and 32 hidden units, followed by the final output layer for prediction. The model was implemented using PyTorch and optimized using the Mean Squared Error (MSE) loss function. Training was conducted using the AdamW optimizer, with a learning rate of 0.01 and a weight decay of ${10}^{-5}$. The validation set is used for hyperparameter optimization, and the test set is used for final model evaluation and performance reporting.

**Machine learning feature importance analysis.** To identify genomic features influencing editing activity, DeepLIFT was used to compute feature importance scores. The on-target sequences served as the baseline input for DeepLIFT. For each position, the scores were summed to quantify the negative impact of having a mismatch at that specific position.

**Feature analysis of variant impact on Cas9 activity.** A variant is defined as a single nucleotide variant occurring at an off-target site that increases the original number of mismatches by 1. For CHANCE-seq data, all such possible variants were enumerated using the BK-tree library <https://github.com/benhoyt/pybktree>. Relative activity was calculated as the ratio of Cas9 off-target enrichment ratio to on-target enrichment. The off-target set at each mismatch number (e.g, mismatch of 2 means an off-target set with all off-targets with mismatch=2 and corresponding variant-containing off-targets with mismatch=3) were used to calculate overall mismatch occurring frequency and transition mismatch frequency. To summarize the high-impact variant effect, we first selected the top 1000 highly active sites for each of the five on-target in CHANCE-seq. Then, the top 100 sites with the most deleterious variant effect were used to draw the off-target context and variant feature heatmap. For CHANCE-net variant feature analysis, a large set of simulated sequences was generated. Specifically, (1) on-target sequences were generated based on the original on-target, to eliminate possibly on-target bases bias at each position. Each new on-target differs to the original on-target by 1 mismatch. The first base G and PAM NGG keep the same. The total number of on-target is 348, including the 6 original on-target sequences used by GUIDE-seq-2. (2) Off-target sequences were generated by randomly mutating X bases, where X equals the predefined number of mismatches. The 21 position N is not counted. This simulation was repeated 10,000 times for each given mismatch. Duplicated off-targets were removed. (3) For each off-target sequence from (2), all possible variant-containing off-targets were subsequently generated by mutating each position to each possible base. The off-target context and variant feature heatmap was generated based on the top 1,000 Cas9-activity sites for each on-target from (1). For each number of mismatches, the number top Cas9-activity sites is 348,000, which was further sorted by variant effect and divided equally into 5 bins (e.g., Quintile 5 represents the most deleterious variant effect).

**Processing of CRISPRoffT data**. The CRISPRoffT dataset was obtained from the publicly available website (<https://ccsm.uth.edu/CRISPRoffT/download.html>). To avoid data leakage, all samples originating from publications co-authored by the authors of this study were removed. Four experimental technologies that measure cellular genome-editing activity, TTISS, PEM-seq, GUIDE-seq, and iGUIDE, were included in the benchmarking analysis. Off-target sites containing more than six mismatches were excluded. For each technology and target, normalized experimental score was calculated as the ratio of off-target activity to on-target activity.

**Evaluation of existing CRISPR-cas9 editing activity models.**

Four state-of-the-art methods were evaluated for comparison: Cutting frequency determination (CFD), CRISPR-Net, CRISPROFF, and CRISOT, all designed to predict Cas9 cutting efficiencies based on an off-target and an on-target sequence. The CFD method is based on a position weight score for each possible mismatch between off-target and on-target. The CRISPR-Net is based on a recurrent convolutional network model. The CRISPROFF method is limited to NGG, NGA, and NAG PAMs. The CRISOT method only works for NGG PAM. In instances where a variant alters the PAM sequence and renders CRISPROFF or CRISOT inapplicable, a cutting efficiency of zero is assumed. To assess the impact of sequence variants, all 4 models were applied to both reference and variant-containing off-targets. Only SNP variants and 23bp off-targets were considered. The difference (i.e, variant effect) in predicted Cas9 cutting efficiencies between the variant and reference sequences was used to quantify the effect of the variant on cleavage efficiency. The high-confidence variants’ effect based on rhAmpSeq indel frequency were used to evaluate these in silico tools against our deep learning method, CHANCE-net. Values for CFD, CRISPR-Net, and CRISOT ranged from 0 to 1 and were multiplied by 100 to be compatible with indel percentages. The variant effect ($\Delta$) was log2 transformed using a symmetrical log function: $sign\left( \Delta\right)*\log_{2} (abs\left( \Delta\right)+1)$. Spearman correlation and Pearson correlation were calculated to evaluate their performance.

**Methods References**
